# The role of the 5’ sensing function of ribonuclease E in cyanobacteria

**DOI:** 10.1101/2023.01.13.523895

**Authors:** Ute A. Hoffmann, Elisabeth Lichtenberg, Said N. Rogh, Raphael Bilger, Viktoria Reimann, Florian Heyl, Rolf Backofen, Claudia Steglich, Wolfgang R. Hess, Annegret Wilde

## Abstract

RNA degradation is crucial for many processes in pro- and eukaryotic organisms. In bacteria, the preference of the central ribonucleases RNase E, RNase J and RNase Y towards 5’-monophosphorylated RNAs is considered important for RNA degradation. For RNase E, the underlying mechanism is termed 5’ sensing. Cyanobacteria, such as *Synechocystis* sp. PCC 6803 (*Synechocystis*), encode RNase E and RNase J homologs. Here, we constructed a *Synechocystis* strain lacking the 5’ sensing function of RNase E and mapped on a transcriptome-wide level 292 5’-sensing-dependent cleavage sites. These included so far unknown targets such as the 5’ untranslated region of the response regulator gene *lsiR*; *trxA, apcE* and *atpI* mRNAs, encoding proteins related to energy metabolism; as well as *sbtB* and *rbcLXS* encoding proteins relevant for carbon fixation. Cyanobacterial 5’ sensing is important for the maturation of rRNA and several tRNAs, including tRNA^Glu^_UUC_. This tRNA activates glutamate for tetrapyrrole biosynthesis in plant chloroplasts and most prokaryotes. We found that increased RNase activities leads to a higher copy number of the major *Synechocystis* plasmids pSYSA and pSYSM. The results provide a first step towards understanding the relative importance of different target mechanisms of RNase E outside *Escherichia coli*.

## Introduction

RNA degradation in bacteria is thought to be initiated by the removal of 5’-pyrophosphates from 5’-triphosphorylated (5’-PPP) RNA ends or by an endonucleolytic cleavage, usually mediated by one or several of the three ribonucleases (RNases) RNase E, RNase J or RNase Y (1, 2). In both cases, the resulting cleavage products are 5’-monophosphorylated (5’-P), which foster further degradation due to the higher affinity of RNase E, RNase J and RNase Y towards 5’-P rather than 5’-PPP RNA ends (1). The mechanism of 5’-monophosphate recognition by RNase E is called 5’ sensing. All three RNases play central roles in RNA degradation in different bacteria (1, 3). For instance, inactivation of RNase E in *Escherichia coli* (*E. coli*) led to a 6-fold increase in the half-life of pulse-labeled mRNA fractions and was ultimately lethal (4). In addition to its role in RNA turnover, *E. coli* RNase E is involved in rRNA (5, 6), tRNA (7–9), tmRNA (10), 6S RNA (11) and sRNA (12–14) maturation, as well as in the targeting of specific mRNAs by certain regulatory sRNAs (12, 15, 16).

On a structural level, the higher affinity of *E. coli* RNase E towards 5’-P RNA fragments is based on a shallow pocket on the protein surface (15, 17) (Fig. 1A). Amino acid exchanges R169Q and T170V reduced the affinity towards 5’-P RNA fragments (6, 15, 18, 19). *In vivo*, the substitution R169Q led to a slow growth phenotype, the accumulation of 5S rRNA precursors and prolonged half-lives of several transcripts (5, 6). The R169Q and T170V substitutions enabled the dissection of the relevance of different target recognition pathways of *E. coli* RNase E (5–8, 19, 20). Besides its high affinity towards 5’-P RNA, there are further determinants directing cleavage by RNase E. The enzyme cleaves single-stranded RNA in A and U rich regions (21). For many transcripts, secondary structures or multiple single-stranded RNA regions in close proximity to the cleavage site are important for processing (7, 8, 15, 22). In absence of secondary structures, a guanine residue two nucleotides (nt) upstream of the cleavage site might be of importance in *E. coli* (23), while in *Salmonella enterica* a uracil residue located two nt downstream directs positioning of the cleavage site (14). However, the interplay among these different cleavage determinants is not yet fully understood and several studies suggested that different target recognition mechanisms can, to a certain extent, substitute for each other (5, 15, 24). There are several targets for which the dependence on 5’ sensing was shown in enterobacteria *in vivo*, e.g. the maturation of the 3’ untranslated region (UTR)-derived sRNA CpxQ (13), sRNA sponge SroC (12) and processing of 9S RNA as well as *rpsT* mRNA (5, 20). *In vitro*, presentation of a 5’-P RNA end is important for MicC-directed cleavage of *ompD* mRNA by RNase E (16). However, a transcriptome-wide survey of processing sites depending on 5’ sensing has not yet been performed in any bacterium *in vivo.* Cyanobacteria are the only bacteria capable of performing oxygenic photosynthesis and are therefore of immense ecological relevance as well as of interest for the phototrophic production of valuable compounds from CO_2_. Cyanobacterial RNA decay is far from being fully understood and potentially important players remain to be characterised experimentally (25). Several studies demonstrated the regulation of photosynthetic transcript accumulation by cyanobacterial RNase E (26–28). In the unicellular model cyanobacterium *Synechocystis* sp. PCC 6803 (*Synechocystis*), RNase E was identified as the major maturation enzyme of crRNAs from a subtype III-Bv CRISPR array (29). The RNase E of cyanobacteria represents a compact version of the enzyme, lacking both the long N-terminal extensions typical for chloroplast RNase E (30), as well as the region homologous to the C-terminal half of enterobacterial RNase E (31). Findings for *E. coli* RNase E and RNase G imply that the C-terminal half of *E. coli* RNase E might be involved in target recognition mechanisms alternative to 5’ sensing (5, 15, 24). Hence, the lack of apparent homologous protein structures might indicate that cyanobacterial RNase E is more dependent on 5’ sensing than its enterobacterial counterpart. Interestingly, cyanobacteria do not only encode RNase E, but also RNase J, which also shows a higher affinity towards 5’-P than to 5’-PPP RNA ends and a similar cleavage affinity as RNase E *in vitro* (32). Recently, we showed that, in contrast to its role in *E. coli*, RNase E activity in *Synechocystis* is not limiting for 5’-dependent RNA degradation (33). Hence, cyanobacteria provide a well-suited model to study the impact of 5’ sensing on cleavage positioning without affecting general 5’-end dependent bulk RNA degradation *in vivo*.

**Figure 1:**
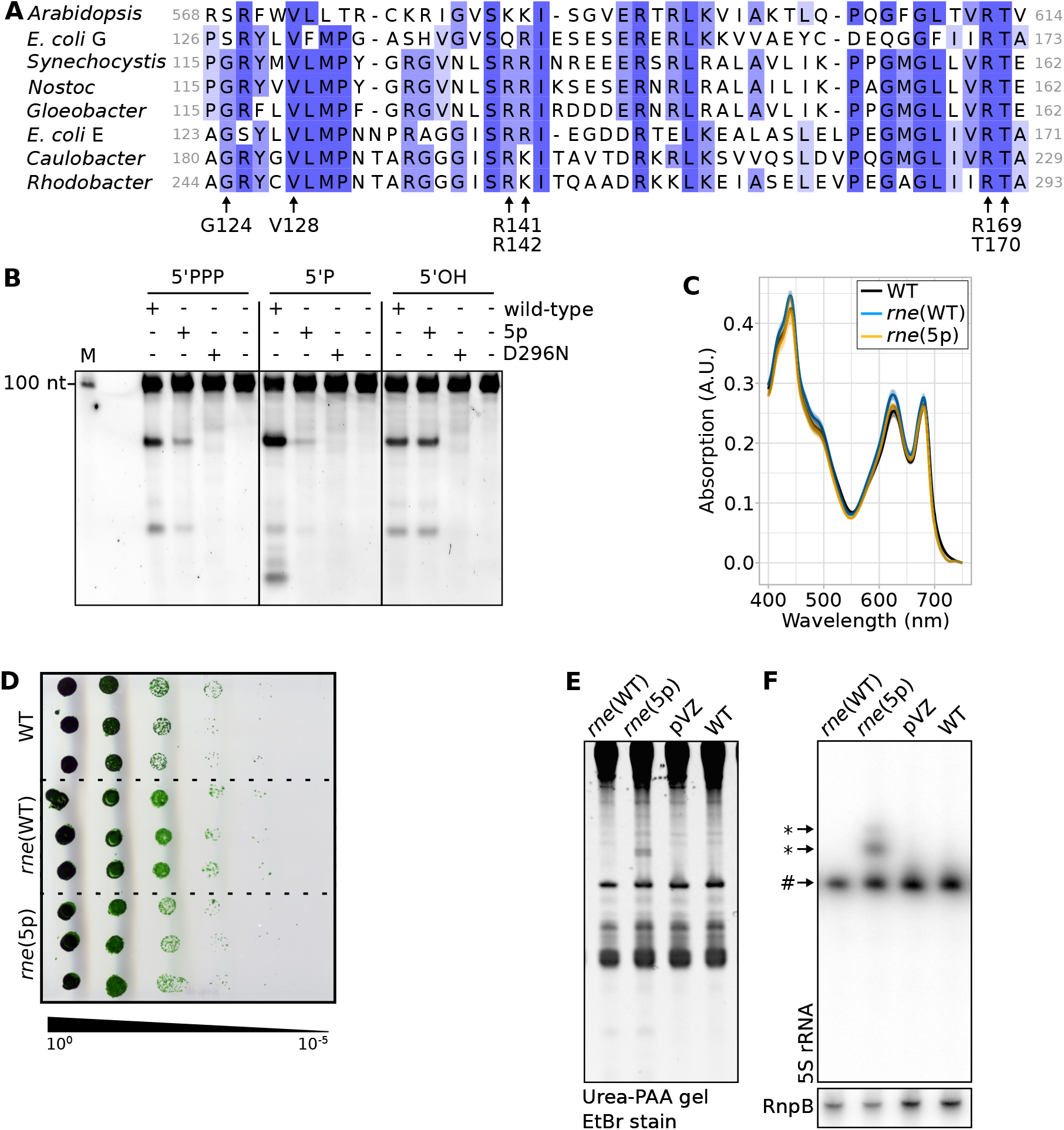
Impact of amino acid exchange T161V and characterisation of *Synechocystis* strains *rne*(WT) and *rne*(5p). (**A**) Alignment of *Synechocystis* RNase E residues 115 to 162 with the respective section in homologues from *Arabidopsis thaliana, Nostoc* sp. PCC 7120, *Gloeobacter violaceus* PCC 7412, *E. coli* (K12), *Caulobacter crescentus* (CB15), *Rhodobacter sphaeroides* 2.4.1 and *E. coli* (K12) RNase G. Arrows point at key amino acids forming the 5’ sensor pocket in *E. coli* RNase E (15, 17). Shades of blue indicate different degrees of identity between the compared protein sequences. (**B**) *In vitro* assay comparing the cleavage activity of *Synechocystis* wild-type RNase E (wild-type), RNase E with amino acid exchange T161V (5p) and catalytically inactive RNase E (29) harbouring the amino acid exchange D296N (D296N) on a 113 nt long transcript derived from the 5’ UTR and the first 20 codons of *psbA2.* The 5’ termini were either triphosphorylated (5’PPP), monophosphorylated (5’P) or hydroxylated (5’OH). One representative SYBR-gold stained 8% urea-PAA gel is shown (n=5). M: marker. (**C**) Absorption spectra of wildtype *Synechocystis* (WT), *rne*(WT) and *rne*(5p). Shaded areas indicate standard deviations of three replicates. (**D**) Growth and viability of WT, *rne*(WT) and *rne*(5p) after 8 days. (**E**) Urea-PAA gel of total RNA of WT, *rne*(WT), *rne*(5p) and empty-vector control strain pVZΔKm^R^ (pVZ) stained with ethidium bromide. (**F**) The same gel after northern blotting and hybridization with a radioactively labelled probe for 5S rRNA. One representative analysis is shown (n=4). # indicates mature 5S rRNA, * 5S rRNA precursors. A control hybridization with RnpB is shown (lower panel).

To elucidate the possible impact of 5’ sensing in cyanobacterial RNase E, we constructed a *Synechocystis* strain with a T161V amino acid exchange, homologous to T170V, which was successfully used in *E. coli* to decrease 5’ sensing activity (8, 19). The resulting strain, *rne*(5p), displayed clear and specific deficiencies in rRNA and tRNA maturation. Furthermore, we identified potential cases of post-transcriptional regulation conveyed by 5’ sensing. Interestingly, many instances of RNase-E-dependent processing were not affected by the decreased 5’ sensing activity, indicating that also the *Synechocystis* enzyme employs further RNA target recognition pathways, despite its shorter C-terminal segments lacking sequence similarity to the *E. coli* enzyme (34).

## Material and Methods

### Overexpression and purification of cyanobacterial RNase E from *E. coli*

A pQE70-based vector containing a codon-optimised version of the *Synechocystis* RNase E encoding *slr1129* gene fused to a C-terminal TEV split site and 6xHis-tag was kindly provided by J. Behler (29). This gene was amplified using primer pair C01/C02 (see supplementary Table S1 for all primer sequences) and subcloned into pJET1.2/blunt (Thermo Fisher Scientific). Primer pairs M01/M02 and M03/M04 were used with the Q5 site-directed mutagenesis kit (NEB) to introduce point mutations for the T161V and D296N substitutions. Constructs encoding wild-type, T161V or D296N RNase E and vector pCDFDuet were cut with the FastDigest restriction enzymes NcoI and SalI (Thermo Fisher Scientific). Insert and vector were ligated using T4 DNA ligase (Thermo Fisher Scientific) and transferred to *E. coli* DH5α. *E. coli* BL21(DE3) was transformed with the resulting plasmids and recombinant *Synechocystis* RNase E was expressed and purified as described previously (35). In short, 50 ml LB medium containing appropriate antibiotics was inoculated with the transformed *E. coli* cells and incubated overnight at 37°C. The next day, this pre-culture was diluted 1:50 in 1,5 L of LB medium containing appropriate antibiotics. This culture was grown to ODß00 = 0.6 to 0.8, as measured using a BioSpectrometer basis (Eppendorf). Protein expression was induced by addition of 1 mM isopropyl-ß-D-thiogalactopyranosid (IPTG) and cultures were further incubated for 16 h at 22°C. Cells were harvested by centrifugation (6,000 g, 10 min, 4°C) and lysed using a French-press (Constant Systems) at 36,000 ps. Cell debris was removed by centrifugation (13,000 g, 30 min, 4°C) followed by filtration through 0.45 μm and 0.22 μm pore sized polyethersulfone membranes (ROTILABO®). RNase E was then purified using TALON® Metal Affinity Resin (Takara Bio). Protein purity was confirmed by SDS-polyacrylamide gel electrophoresis (PAGE) followed by Coomassie staining. Enzymatic activity of RNase E was analyzed by *in vitro* cleavage assays using 5’PPP and 5’P *psbA2* transcripts (35).

### Generation of transcripts with different 5’ modifications for *in vitro* cleavage assays

PCR products for the generation of different potential RNase E targets were amplified with MyTaq™ Red Mix 2x (Bioline), and *Synechocystis* genomic DNA as template (primers T01 to T12). Substrates were transcribed *in vitro* from purified PCR products using the HiScribe™ T7 Quick High Yield RNA Synthesis Kit (NEB) according to manufacturer’s instructions. For the 53 nt *atpT* substrate, the two primers were annealed to each other and used as *in vitro* transcription template. To remove DNA templates, transcripts were treated with 4 U DNase I (NEB) for 30 min at 37°C and subsequently separated on urea-polyacrylamide (PAA) gels (8.3 M urea, 10% PAA). Correctly sized transcripts were excised and purified using the ZR small-RNA™ PAGE Recovery kit (Zymo Research). Buffers were exchanged using RNA clean & concentrator-25 kit (Zymo Research), eluting RNA in RNase-free H_2_O. For differently modified 5’termini, various treatments were performed. To obtain 5’P, 2 μg 5’PPP transcripts were incubated with 10 U RppH (NEB) and 20 U RiboLock (Thermo Fisher Scientific) in NEBuffer 2 (NEB) for 2 h at 37°C. Successful RppH treatment was verified by incubating 100 ng of 5’P or 5’PPP transcripts with 20 U RiboLock and 1 U Terminator™ 5’Phosphate-Dependent Exonuclease (Lucigen) in 1x TEX Buffer B for 30 min at 42°C. Exemplary results for four transcripts are shown in supplementary Figure S1. To obtain 5’-hydroxylated termini, transcripts were treated with Shrimp Alkaline Phosphatase (SAP; from NEB). In short, 2 μg RNA were mixed with 10 U RiboLock, 3 U SAP in 1x SAP buffer and incubated for 2 h at 37°C. After these treatments, enzymes were removed and buffers exchanged using the RNA clean & concentrator-25 kit (Zymo Research). Modified transcripts were eluted in 15 μl H_2_O.

### RNase E *in vitro* cleavage assays

*In vitro* cleavage assays were performed according to (29, 35). Briefly, 80 ng RNA substrates (5’PPP, 5’P, 5’OH) were treated in 5 μl or 10 μl reaction volume containing RNase E cleavage buffer (25 mM Tris-HCl, 60 mM KCl, 5 mM MgCl_2_, 100 mM NH_4_Cl, 0.1 mM DTT, pH 8.0) and 1 μl or 2 μl RNase E eluates (wild-type, T161V, D296N) respectively. The reactions were performed at 30°C for 30 min and stopped by adding 2x loading dye (0.025% xylene cyanol, 0.025% bromophenol blue, 0.5 mM EDTA, 0.025% SDS, and 95% formamide). Transcripts were denatured for 10 min at 65°C and depending on their size loaded on 6%, 8% or 15% 8.3 M urea-PAA gels. Gels were stained by incubation in 1x SYBR Gold Nucleic Acid Gel Stain (Invitrogen) in 0.5x Tris-borate-EDTA buffer for 15 min. Signals were detected using a Typhoon™ FLA 9500 (Cytiva) imager, with following settings: excitation light wavelength: 495 nm; emission light wavelength: 537 nm; emission filter: long pass blue (LPB); photomultiplier value: 450 – 600 V.

### Bacterial strains and culture conditions

We used a motile wild type of *Synechocystis*, which is capable of photoautotrophic, mixotrophic and chemoheterotrophic growth on glucose. Originally, it was obtained from S. Shestakov (Moscow State University, Russia) in 1993 and re-sequenced in 2012 (36). For culturing, BG-11 medium (37) substituted with 0.3% (w/v) sodium thiosulfate and 10 mM N-[Tris-(hydroxy-methyl)-methyl]-2-aminoethanesulfonic acid (TES) buffer (pH 8.0) was used. Liquid cultures were grown in Erlenmeyer flasks under constant shaking (135 rpm) at 30°C and continuous white-light conditions (30 μmol photons m^-2^ s^-1^). Plate cultures were grown on 0.75% bacto-agar BG-11 plates. Kanamycin (40 μg ml^-1^) and chloramphenicol (10 μg ml^-1^) were added to plate cultures for strain maintenance, but omitted during experiments.

### Construction of mutant strains

Information on all used strains is summarised in supplementary Table S2. Strain *rne*(5p) was constructed as described previously for *rne*(WT) (33) (Supp. Fig. S2A). Primers M03/M04 (Supp. Table S1) were used for introducing the T161V mutation into the pJET 1.2 vector harbouring the *rne-rnhB* operon including N-terminal 3xFLAG tag, promoter and terminator sequences (33).

### Spectroscopy

Whole-cell absorption spectra were measured using a Specord 250 Plus (Analytik Jena) spectrophotometer at room temperature and normalised to absorption values at 750 nm which were measured with a BioSpectrometer basic (Eppendorf).

### Spot assays

Spot assays were performed as described previously (38). Plates were incubated at continuous white light (50 μmol photons m^-2^ s^-1^).

### DNA extraction

To extract DNA for homozygosity testing, 0.2% glucose was added 12 h prior to harvesting 50 ml cultures by centrifugation (6000 g, 10 min, 4°C). To test relative replicon abundances, DNA was extracted from 50 ml culture at OD_750nm_ 0.6 to 0.7 without prior glucose addition. After centrifugation, cell pellets were washed twice using 10 mM Tris/HCl, 1 mM EDTA buffer (pH 8.0), resuspended in 1 ml TES-buffer (25% (w/v) sucrose, 100 mM EDTA, 50 mM Tris/HCl pH 8.0) and snap-frozen in liquid nitrogen. Samples were thawed at 60°C and again snap-frozen in liquid nitrogen. After thawing, lysozyme (5 mg ml^-1^) and RNase A (0.1 μg ml^-1^) were added and samples were incubated at 37°C for 1 h. Proteinase K (40 μg ml^-1^) and SDS (2%) were added and samples were incubated for at least 4 h at 60°C. This was followed by two extractions using phenol-chloroform-isoamyl alcohol (25:24:1 (v/v)) and one extraction using chloroform. DNA was precipitated from the aqueous phase with 0.7 vol. isopropanol overnight. After centrifugation, pellets were washed with ethanol (70%). After residual traces of ethanol were evaporated, pellets were resuspended in water.

### Southern blot analysis

To test full segregation of *rne*(WT) and *rne*(5p), restriction digests of 5 μg of DNA extracted from three independent clones were performed with HincII or Bsu15I (Thermo Fisher Scientific). Duplicates of wild-type *Synechocystis* (WT) DNA were included as control. To test copy numbers of different replicons, 5 μg of DNA extracted from three independent liquid cultures of WT, pVZΔKm^R^ and *rne*(5p) and four *rne*(WT) cultures was cleaved using enzymes HindIII and PstI (Thermo Fisher Scientific). Samples were mixed with DNA loading dye (4.5% (v/v) glycerol, 0.05% bromophenol blue, 0.05% xylene cyanol, 0.05% orange G, Midori Green) and separated by gel electrophoresis on 0.8% (w/v) agarose gels in Tris-Acetate-EDTA buffer (40 mM Tris, 1 mM EDTA pH 8.0, 20 mM acetic acid). Subsequently, gels were incubated in 0.25 M HCl until 15 min after the bromophenol blue band in the gel turned green. After washing with ddH_2_O, gels were incubated in 1.5 M NaCl, 0.5 M NaOH until 15 min after the bromophenol blue band turned blue again. Gels were washed with ddH_2_O and incubated in 1.5 M NaCl, 0.5 M Tris/HCl pH 7.5 for 20 min. DNA was transferred onto Roti-Nylon plus membranes (Carl Roth) by capillary blotting. To test full segregation of strains by Southern blot hybridization, a probe was labeled using the Amersham Gene Images AlkPhos Direct Labelling and Detection System (GE Healthcare) according to the manufacturer’s protocol with a DNA fragment generated by PCR using primer pair S01/S02 (Supp. Fig. S2A). Luminescence signal was detected with the Fusion SL4 advanced (Vilber Lourmat) apparatus. To test replicon abundance, membranes were hybridised with radioactively labelled RNA probes and signals were detected as described for northern blot analysis (33). Primer pairs S03/S04 (chromosome), S05/S06 (pSYSA) and S07/S08 (pSYSM) were used to generate DNA templates for probe transcription. Signal intensity was quantified with Fiji (v2.0.0-rc-69/1.52p) (39), followed by analysis using an R (v4.0.4) script. For statistical analysis, ANOVA followed by post-hoc Tukey’s honest significant difference tests were performed.

### Northern blot hybridization

For RNA extraction from *Synechocystis*, liquid cultures were grown to OD_750nm_ of 0.6 to 0.8. Sampling of 20 ml or 30 ml aliquots and RNA extraction and northern blot hybridization were performed as described (33). Primer pairs N01/N02 and S05/S06 were used to generate DNA templates for *in vitro* transcription and hybridization as described (33). To create the probe complementary to the 5’ end of tRNA^Glu^_UUC_, oligos N03 and N04 were annealed by mixing 400 pmol of each in 8 μl ddH2O. This mixture was heated to 95°C and then cooled down to 20°C with a stepwise decrease of 0.1°C per second. The annealed oligos were used for *in vitro* transcription. Hybridization was performed at 55°C and washing at 50°C. Oligonucleotide 5S-rRNA-oligo was labelled using T4 PNK (NEB) and [γ-32-P]-ATP (Hartmann Analytics, Germany) and hybridization was carried out as described (29, 33) except that formamide was omitted from the hybridization buffer. Hybridization was performed at 45°C overnight. After 10 min incubation with washing solution I (2x SSC, 1% SDS), membranes were incubated at 45°C for 10 min each with washing solutions I and subsequently II (1x SSC, 0.5% SDS).

### RNA sampling, extraction and sequencing

For RNA-sequencing, three independent 50 ml liquid cultures of WT, *rne*(WT) and *rne*(5p) were grown to an OD_750nm_ of 0.7 to 0.8. Aliquots of 30 ml were sampled. Small aliquots of culture were tested by PCR for full segregation of mutant strains as described previously (33). One replicate of *rne*(5p) was not fully segregated and excluded from further procedures. After RNA extraction, DNase treatment, subsequent RNA recovery and RNA integrity control were performed as described (33). cDNA libraries were constructed and sequenced as a service provided by Vertis Biotechnologie AG (Germany) according to the tagRNA-Seq protocol (40). Library preparation was performed as described previously (33) with the following modifications: After ligation of processing site (PSS)-specific sequencing adaptors, unligated 5’-P RNA fragments were removed using terminator exonuclease (TEX, Lucigen), followed by ligation of transcriptional start site (TSS)-specific adaptors.

### Bioinformatic analyses

RNA-Seq analysis was performed as described (33) with several modifications detailed below. Reads were uploaded to the public usegalaxy.eu server and analysed utilizing the Galaxy web platform (41) after preliminary processing. Here we used the *Synechocystis* genome annotation as introduced by Kaneko et al. (42) with *sll/slr* and *ssl/ssr* gene identifiers for longer and shorter protein-coding genes and *ncl/ncr* identifiers for non-coding RNAs according to Kopf et al. (43). If gene functions were assigned, we use the respective specific gene names. Several workflows were created to process the data further. These can be accessed and reproduced at the links given in the data availability statement. Subsequently, htseq-count files, PSS/TSS-5’ end files and transcript coverage files were downloaded and analysed using Python (v3.7.4) and R (v4.0.4) scripts available at (https://github.com/ute-hoffmann/5sensing_Synechocystis, Zenodo repository https://doi.org/10.5281/zenodo.7509658). Gene set enrichment analysis (GSEA) (44) was performed using clusterProfiler (45). For classification of genomic positions as TSS or PSS, DESeq2 (46) was used as described (33). PSS positions identified previously (33) were included in subsequent analyses. The resulting set of PSS and TSS positions were used for differential expression analysis with DESeq2 (|log_2_FC| > 1.0 and p.adj < 0.05). Reads with a mapping quality of exactly one, which correspond to multi-mapping reads, were included for the analysis of rRNA loci. Sequence logos were created using WebLogo (v3.7.8) (47, 48) with a GC content of 47.4%. RNA fold (v2.4.17) (49) was used to calculate the minimal folding energy (ΔG) at 30°C and using a sliding window approach with a 1 nt step size. The data was averaged among replicates using an R script and visualised with Artemis (50–52). Figures were created using Inkscape (v1.0.2). The sequence alignment was conducted with MUSCLE (53, 54) and analysed using Jalview (v2.11.0) (55).

## Results

### Amino acid exchange T161V abolishes 5’ sensing of *Synechocystis* RNase E *in vitro*

Recently, we found that *Synechocystis* RNase E activity is not limiting for 5’-dependent RNA degradation *in vivo* (33), even though amino acids which were shown to be important for 5’ sensing in *E. coli* are highly conserved among RNase E homologues (G124, V128, R141, R142, R169, T170; Fig. 1A). At the same time, compared to *E. coli* RNase E, the *Synechocystis* enzyme lacks the extensive C-terminal domain, which seems to be important for target recognition pathways alternative to 5’ sensing in *E. coli* (5, 15, 24). Hence, we asked what is the role and importance of RNase E 5’ sensing for *Synechocystis*? To answer this question, we introduced the amino acid exchange T161V, which is homologous to T170V used in *E. coli* to abolish 5’ sensing (Fig. 1A) (8, 19). *In vitro* cleavage assays on a well-established target of *Synechocystis* RNase E, the 5’ UTR of *psbA2*, illustrated that this exchange did indeed abolish the enzyme’s capability to distinguish between different 5’ phosphorylation states (Fig. 1B). Whereas wild-type RNase E exerted much stronger activity on 5’-P *psbA2* than on the 5’-PPP or 5’-hydroxylated (5’-OH) counterparts, the T161V mutant enzyme showed the least activity towards the 5’-P substrate. Moreover, the smallest prominent band was not produced by the T161V mutant enzyme at all, indicating that 5’ sensing is important for positioning cleavage sites *in vitro.* Incubation with the catalytically inactive D296N RNase E mutant enzyme served as a negative control and to test for contaminating RNases in the enzyme purifications. Compared to RNA incubated with the catalytically active RNase E enzymes, only minor degradation was observable after incubation with the catalytically inactive RNase E mutant enzyme.

### T161V-substituted RNase E can replace the wild-type enzyme *in vivo* if expressed from a plasmid

Based on these *in vitro* results, we set out to investigate the importance of 5’ sensing *in vivo*. However, manipulating RNase E in *Synechocystis* poses several challenges. RNase E is encoded by the essential gene *rne* in a dicistronic operon together with *rnhB*, which encodes RNase HII (33). Evidence is accumulating that transcription of this operon is autoregulated by RNase E (32, 33, 56). Furthermore, *Synechocystis* is polyploid. Since RNase E likely also forms a homotetramer in *Synechocystis*, as in *E. coli*, it is crucial to obtain fully segregated clones since wild-type monomers can rescue mutations in other monomers of a tetramer (57). We aimed at comparing strains encoding either wild-type RNase E or RNase E with the amino acid exchange T161V. To engineer strains, we introduced the *rne-rnhB* operon including 3xFLAG tag, promoter and terminator sequences either at a putative neutral site on the chromosome or on a conjugative, self-replicating plasmid. In a second step, we attempted to delete the original *rne-rnhB* locus (Supp. Fig. S2A). In the following, strains for which the plasmid-based strategy was used will be referred to as *rne*(WT) and *rne*(5p). In *rne*(5p), *rne* carries the T161V mutation. The full deletion of the endogenous *rne-rnhB* locus, i.e. full segregation (homozygosity), was reached once the respective plasmids had been introduced (Supp. Fig. S2B). As a control to *rne*(WT) and *rne*(5p), strain pVZΔKm^R^ was constructed, in which the same plasmid backbone without the *rne-rnhB* operon, was introduced. In a second step, a kanamycin resistance cassette was introduced chromosomally upstream of the *rne-rnhB* locus.

Introducing the operon on the RSF1010-based plasmid probably led to a higher gene dosage (58), which, in turn, led to an elevated *rne-rnhB* transcript level (Supp. Fig. S2C, compare (33)). At standard growth conditions, *rne*(WT) and *rne*(5p) did not show a strong obvious phenotype compared to WT (Fig. 1C and D). On RNA level, we observed an accumulation of 5S rRNA precursors in *rne*(5p) (Fig. 1E and F).

### Inactivating 5’ sensing impacts RNA processing and thereby enables identification of unknown targets of cyanobacterial RNase E

To investigate effects of *rne-rnhB* overexpression and the amino acid exchange T161V on a transcriptome-wide level, we performed RNA-sequencing of triplicates of WT, *rne*(WT) and duplicates of *rne*(5p) (Fig. 2A). cDNA libraries were prepared according to the tagRNA-seq protocol (40) (Fig. 2B). tagRNA-seq enables to distinguish 5’-P RNA species, which originate from processing and are thus PSS, and 5’-PPP RNA species, which are typical TSS. In total, we obtained 95.5 million reads for 8 samples. Of these, we mapped 82.2% with high quality to the *Synechocystis* genome. This corresponded to, on average, 9.8 million reads per sample, of which 3.8% were classified as being associated with PSS, 19.1% with TSS, while 77.1% were tagged neither as PSS nor TSS and give general transcript coverage (Supp. Fig. S3A).

**Figure 2:**
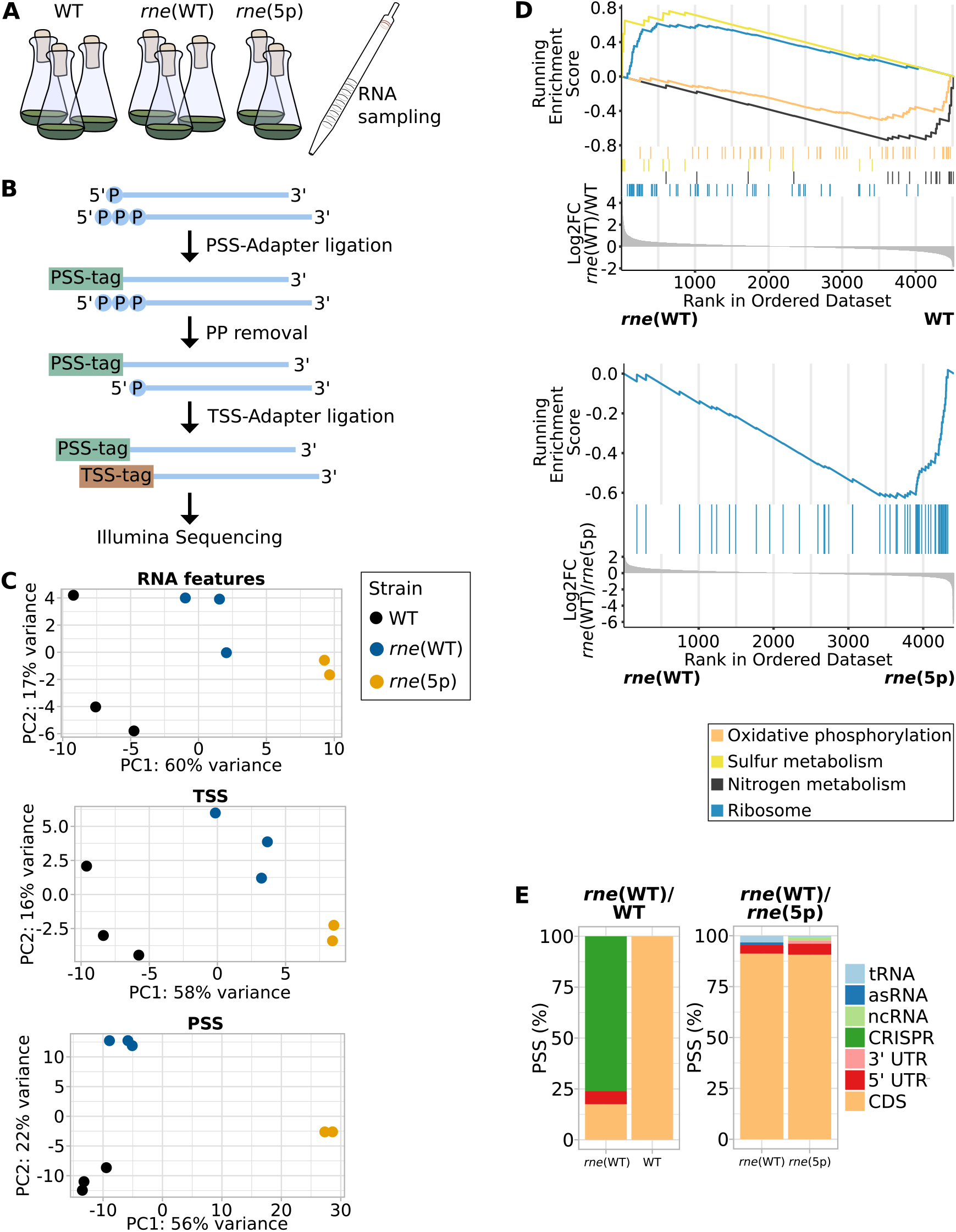
Overview of the RNA-seq experiment. (**A**) Experimental set-up. RNA was sampled from triplicates of wild-type *Synechocystis* (WT), *rne*(WT) and duplicates of *rne*(5p). (**B**) RNA was extracted and used for cDNA library preparation distinguishing between processing sites (PSS), transcriptional start sites (TSS) and other (unspecified) transcript positions. PSS- and TSS-tags, which are sequencing adaptors containing PSS- and TSS-specific nucleotide combinations, were ligated to the respective RNA 5’ ends before or after TEX treatment and 5’ pyrophosphate removal. (**C**) Principal component analyses comparing replicates of the different strains for reads associated with TSS, PSS and other transcript positions. The latter was analysed on RNA features level. (**D**) Gene set enrichment analysis (GSEA) using KEGG terms and unspecified transcript data analysed on level of different RNA features. (**E**) Percentages of different types of RNA features associated with PSS identified in the comparison of *rne*(WT) and WT (on the left) and the comparison of *rne*(WT) and *rne*(5p) (on the right).

As already observed upon transient inactivation of RNase E (33), neither the ratio of PSS to TSS reads nor the ratio of PSS to TSS counts at TSS positions were affected by *rne-rnhB* overexpression or decreased 5’ sensing activity (Supp. Fig. S3). Principal component analyses illustrated consistency among replicates and that the amino acid exchange T161V introduced in *rne*(5p) mainly affected processing (Fig. 2C). This was also reflected by the number of differentially expressed transcripts, TSS and PSS between the three different strains (Table 1 and 2). In total, we found 292 5’-sensing-dependent PSS (subsets of most strongly affected PSS and transcripts in Tables 3, 4 and full list of PSS in Supp. Tables S5 to S12). Many of these PSS overlapped with RNase-E-dependent PSS that we identified recently using the TIER-seq approach (Supp. Fig. S4, Supp. Results S1) (33). We also identified so far unknown targets of RNase E, and 5’-sensing dependent PSS, such as in *trxA* encoding thioredoxin, *apcE* encoding a phycobilisome core-membrane linker, *atpI* encoding ATP synthase A chain of CF(0), *sbtB* encoding P_II_-like signalling protein SbtB, *rbcLXS* encoding Rubisco large and small subunits as well as Rubisco chaperone RbcX and *grpE*, encoding the nucleotide exchange factor GrpE. These transcripts encode proteins with central functions in cyanobacterial metabolism.

**Table 1:** Number of RNA features, transcriptional units (TUs), processing sites (PSS) and transcriptional start sites (TSS) differentially expressed between *rne*(WT), *rne*(5p) and wild-type *Synechocystis* (WT). RNA features include coding sequences (CDS), known sRNAs, asRNAs and small proteins (42, 101, 102) as well as 5’ untranslated regions (UTRs) and 3’ UTRs based on the total of 4,091 TUs defined by Kopf et al. (43). PSS and TSS include positions identified previously (33). Numbers in parentheses give percentages relative to all features of the respective type.

|  | Total | *rne*(WT)/WT |  | *rne*(WT)/*rne*(5p) |  |
| --- | --- | --- | --- | --- | --- |
|  |  | WT | *rne*(WT) | *rne*(5p) | *rne*(WT) |
| RNA features | 6,117 | 76 (1.2%) | 87 (1.4%) | 36 (0.6%) | 32 (0.5%) |
| TUs | 4,091 | 52 (1.3%) | 106 (2.6%) | 30 (0.7%) | 29 (0.7%) |
| TSS | 4,440 | 77 (1.7%) | 83 (1.9%) | 18 (0.4%) | 14 (0.3%) |
| PSS | 6,490 | 7 (0.1%) | 73 (1.1%) | 187 (2.7%) | 105 (1.6%) |

**Table 2:** Number of RNA features differentially expressed between *rne*(WT), *rne*(5p) and WT. Numbers in parentheses give percentages relative to all RNA regions of respective type. “Total” gives the total number of features of the respective type.

|  | Total | *rne*(WT)/WT |  | *rne*(WT)/*rne*(5p) |  |
| --- | --- | --- | --- | --- | --- |
|  |  | WT | *rne*(WT) | *rne*(5p) | *rne*(WT) |
| CDS | 3,675 | 54 (1.5%) | 72 (2.0%) | 26 (0.7%) | 13 (0.4%) |
| 5' UTR | 979 | 6 (0.6%) | 4 (0.4%) | 1 (0.1%) | 0 (0.0%) |
| 3' UTR | 29 | 0 (0.0%) | 0 (0.0%) | 0 (0.0%) | 2 (6.9%) |
| tRNA | 42 | 0 (0.0%) | 0 (0.0%) | 0 (0.0%) | 0 (0.0%) |
| asRNA | 1,071 | 5 (0.5%) | 5 (0.5%) | 2 (0.2%) | 9 (0.8%) |
| ncRNA | 318 | 11 (3.5%) | 3 (0.9%) | 5 (1.6%) | 7 (2.2%) |
| CRISPR | 3 | 0 (0.0%) | 0 (0.0%) | 0 (0.0%) | 1 (33.3%) |

**Table 3:**
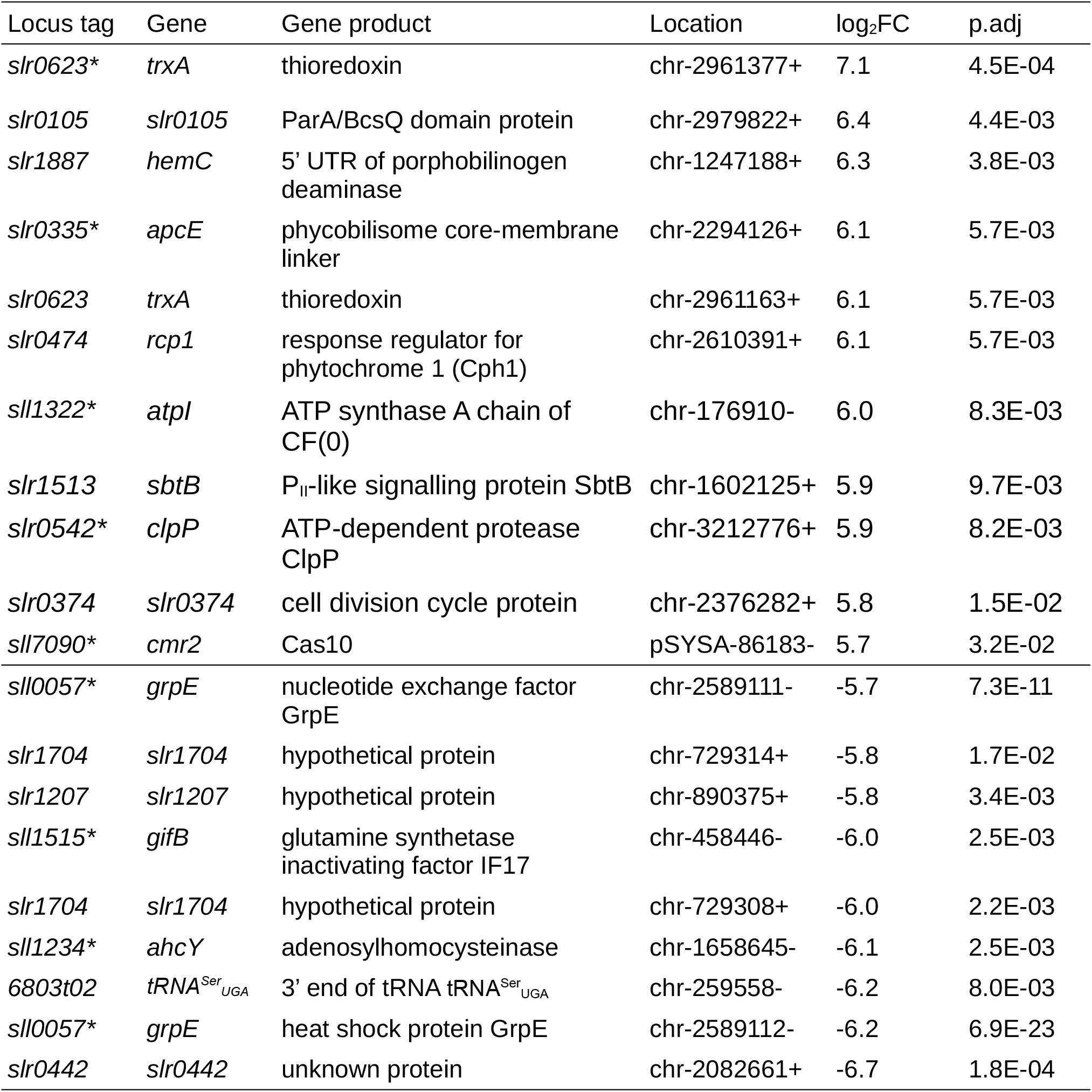
Subset of processing sites (PSS) differentially regulated between *rne*(WT) and *rne*(5p) with |log_2_FC|>5.7 and defined adjusted p values. PSS are sorted according to 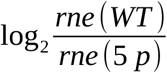. Location is given as (replicon)-(nucleotide position)(strand). chr: chromosome. p.adj: adjusted p value. PSS marked with an asterisk (*) were also identified by TIER-seq to be RNase-E-dependent (33). A full list of PSS with differential regulation is given in Supp. Table S10.

**Table 4:** Subset of RNA features differentially regulated between *rne*(WT) and *rne*(5p) with |log_2_FC| > 1.3 and defined adjusted p values. Rows are sorted according to 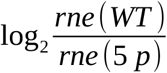.

| Locus tag | Gene | Gene product | $\log_2\text{FC}$ | p.adj |
| --- | --- | --- | --- | --- |
| *sll0327* | *sll0327* | unknown protein | 1.9 | 5.6E-03 |
| *CRISPR2* | *CRISPR2*<br>array | CRISPR2 array | 1.8 | 7.0E-04 |
| *slr0744-as2* | *slr0744-as2* | asRNA to *infB* (translation initiation factor IF-2) | 1.6 | 9.8E-06 |
| *slr1176-as2* | *slr1176-as2* | asRNA to *glgC* (glucose-1-phosphate adenylyltransferase) | 1.6 | 4.8E-02 |
| *sll0247-as2* | *isrR* | asRNA IsrR to *isiA* (iron-stress induced chlorophyll-binding protein) | 1.5 | 5.0E-06 |
| *sll0754-as* | *sll0754-as* | asRNA to *rbfA* (Ribosome-binding factor A) | 1.5 | 3.3E-02 |
| *ncr1010* | *ncr1010* | Ncr1010 | 1.5 | 4.9E-02 |
| *ssl1498* | *ssl1498* | hypothetical protein | 1.4 | 2.4E-02 |
| *sll1671* | *sll1671* | hypothetical protein | 1.4 | 3.2E-02 |
| *ncl0170* | *ncl0170* | Ncl0170 | 1.4 | 1.5E-03 |
| *ncr0710* | *ncr0710* | Ncr0710 | 1.3 | 3.3E-02 |
| *slr0993-as2* | *slr0993-as2* | asRNA to *nlpD* (lipoprotein NlpD) | 1.3 | 4.8E-02 |
| *ncl0350* | *ncl0350* | Ncl0350 | 1.3 | 1.7E-02 |
| *slr2017* | *pilA11* | Minor pilin | -1.3 | 3.7E-02 |
| *sll0944-as* | *sll0944-as* | asRNA | -1.3 | 6.4E-03 |
| *slr1259* | *slr1259* | hypothetical protein | -1.3 | 6.6E-03 |
| *sll1515* | *gifB* | glutamine synthetase inactivating factor IF17 | -1.3 | 5.0E-06 |
| *slr5054* | *xssM* | glycosyltransferase essential for synechan production | -1.3 | 1.3E-05 |
| *slr1544* | *slr1544* | unknown protein | -1.4 | 3.2E-02 |
| *sll1483* | *sll1483* | periplasmic protein | -1.4 | 4.3E-04 |
| *ssr1966* | *ssr1966* | hypothetical protein | -1.4 | 3.4E-03 |
| *sll1514* | *hspA* | 16.6 kDa small heat shock protein | -1.6 | 9.7E-04 |
| *ncl0710* | *ncl0710* | Ncl0710 | -1.6 | 1.7E-04 |
| *sll1862* | *sll1862* | unknown protein | -1.6 | 1.8E-04 |
| *sll1863* | *sll1863* | unknown protein | -1.7 | 3.8E-03 |
| *slr0442* | *slr0442* | 5'UTR of unknown protein | -1.8 | 6.6E-03 |
| *sll0330* | *fabG* | sepiapterine reductase | -1.9 | 3.4E-03 |
| *ssr2848* | *ssr2848* | unknown protein | -2.0 | 1.2E-03 |
| *ncr1420* | *ncr1420* | Ncr1420 | -2.2 | 1.4E-05 |
| *ncl0570* | *ncl0570* | Ncl0570 | -2.8 | 6.1E-07 |
| *ncl1700* | *ncl1700* | Ncl1700 | -4.5 | 8.2E-17 |
| *6803r06* | *rrn5Sb* | 5S rRNA | -5.4 | 2.2E-50 |
| *ncr1230* | *ncr1230* | Ncr1230 | -6.0 | 1.2E-27 |
| *6803r01* | *rrn5Sa* | 5S rRNA | -6.2 | 8.3E-61 |

Both sets of PSS with higher counts in *rne*(WT) or *rne*(5p) showed a strong similarity to the sequence logo identified for *Synechocystis* RNase E previously (33) (Supp. Fig. S5A, B). PSS enriched in *rne*(WT) showed a preference towards single-stranded regions (Supp. Fig. S5C). Due to the low number of PSS differing between *rne*(WT) and WT, we did not create sequence logos for this set of PSS.

### Overexpression of the *rne-rnhB* operon led to increased copy numbers of pSYSA and pSYSM plasmids

Our successful mutant construction strategy resulted in an overexpression of the *rne-rnhB* operon in *rne*(WT) and *rne*(5p) compared to WT. We evaluated how the overexpression of the encoded two RNases, RNase E and RNase HII, affected the transcriptome. Gene set enrichment analyses (GSEA) using KEGG terms revealed that transcripts related to several biological processes, such as oxidative phosphorylation or nitrogen metabolism, were affected in *rne*(WT) compared to WT (Fig. 2D, Supp. Tables S3 and S4). Compared to the effects observed upon transient inactivation of *Synechocystis* RNase E (33), the overexpression of the *rne-rnhB* operon in *rne*(WT) compared to WT had only a relatively mild effect on the transcriptome.

The majority of PSS with higher counts in *rne*(WT) than in WT corresponded to processing events in the CRISPR3 array (64.4%, Fig. 2E). This is consistent with prior findings comparing an RNase E overexpressing strain to WT (29). Accordingly, mature crRNAs of the CRISPR3 array accumulated strongly in *rne*(WT) and *rne*(5p) compared to WT and the empty-vector control strain pVZΔKm^R^ (Supp. Fig. S6). Further PSS with a higher read count in *rne*(WT) than in WT mapped to the 5’ UTR and coding sequence (CDS) of *rne*, 5’ UTRs of *gifA* and *pirA* (*ssr0692*) as well as the CDS of *gifB.*

In our RNA-seq analysis, we observed a general upregulation of transcripts encoded on plasmids pSYSA (padj=2*10^-10^), pSYSM (padj=2*10^-10^) and pSYSX (padj=2.1*10^-7^) in *rne*(WT) compared to WT (Fig. 3A, left side), suggesting that overexpression of the *rne-rnhB* operon affected transcription of plasmid-located genes more generally. This effect was more pronounced in *rne*(5p) for pSYSM (padj=2.2*10^-8^) and in *rne*(WT) for pSYSA (padj=4*10^-10^) (Fig. 3A, right side). pSYSX transcript levels were not influenced significantly by the inactivation of 5’ sensing. Transcript level differences between different strains were not reflected by the distribution of PSS to the five plasmids (Supp. Fig. S7). Hence, we assumed these differences were not based on a general preference of RNase E to process plasmid- or chromosome-derived transcripts. Alternatively, increased plasmid copy numbers may lead to a generally higher level of plasmid-encoded transcripts. To test for this, we performed Southern blot and qPCR analyses (Fig. 3B, C and Supp. Fig. S8). Indeed, we found that the pSYSA hybridization signal was significantly higher in *rne*(5p) (~2.4 fold compared to WT) and *rne*(WT) (~3.8 fold compared to WT) compared to WT and the empty-vector control strain pVZΔKm^R^ (padj < 0.05). For pSYSM, this was the case for *rne*(WT) compared to WT and pVZΔKm^R^ (~2.4 fold higher compared to WT).

**Figure 3:**
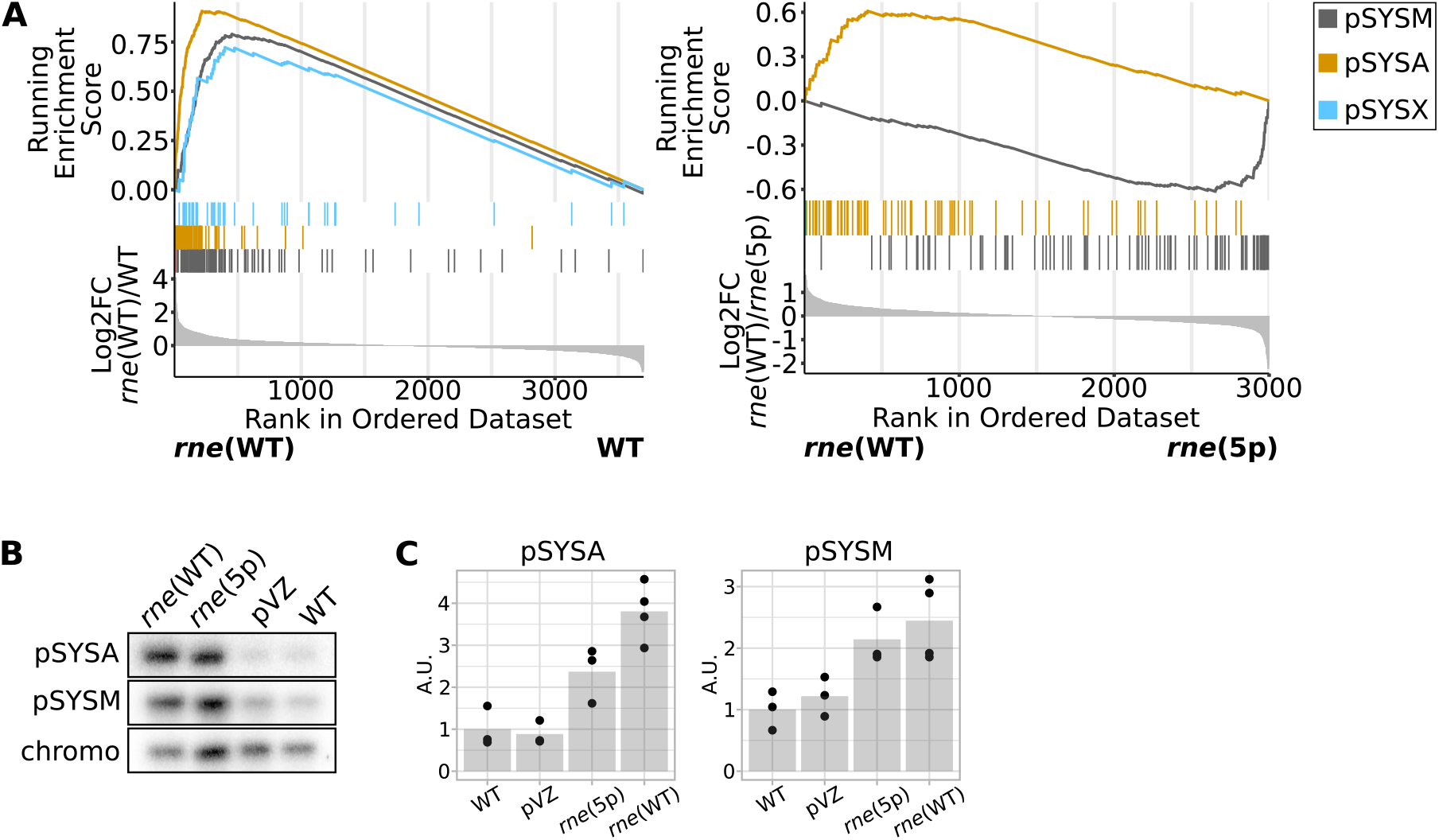
Influence of overexpression of *rne-rnhB* operon on transcriptome and copy number of plasmids pSYSA and pSYSM. (**A**) Gene set enrichment analysis (GSEA) comparing enrichment of transcripts encoded on different replicons. Data is only shown for replicons with significant differences between strains. (**B**) Southern blot comparing relative amounts of pSYSA, pSYSM and chromosomal DNA in different strains. DNA was digested with HindIII restriction enzyme. One representative analysis is shown (n=3). chromo: hybridization signal corresponding to chromosomal DNA. (**C**) Quantification of hybridization signals for pSYSA and pSYSM, normalised by chromosomal hybridization signal. Dots represent biological replicates, gray bars the means of replicates, normalised to the WT values. pVZ: empty-vector control strain pVZΔKm^R^.

### rRNA and tRNA maturation as well as transcript levels of several ribosomal proteins are dependent on RNase E 5’ sensing

Compared to other RNA feature types, a high proportion of PSS differing between *rne*(WT) and *rne*(5p) mapped to tRNAs (Fig. 2E and 4A). Peaks included positions both with a higher and with a lower read count in *rne*(5p) compared to *rne*(WT) (Table 5). These PSS mapped both 5’ and 3’ of the mature tRNAs and in two instances (tRNA^Ser^_UGA_ and tRNA^Asp^_GUC_) exactly 1 nt downstream of the mature 3’ end (Supp. Fig. S9A). Several of these RNase-E-dependent PSS were located precisely at the 5’ ends of mature tRNAs (Fig. 4B). We mapped a 5’-sensing-dependent PSS to the *6803t14* locus, which encodes the tRNA^Glu^_UUC_ (Fig. 4B). This glutamyl-tRNA is of interest beyond its role in translation because it activates glutamate as the first committed precursor for tetrapyrrole biosynthesis in plants, cyanobacteria and many other bacteria as well as Archaea (59–61). The PSS peak, which is located at the mature tRNA’s 5’ end, is clearly reduced in *rne*(5p) compared to *rne*(WT) (log_2_FC(*rne*(WT)/*rne*(5p) = 1.6, padj=0.015, Fig. 4B, Table 5). A similar reduction was observed when RNase E activity was lowered in the strain with the temperature-sensitive RNase E (Supp. Fig. S9B). Additionally, we tested the effect in strain *rne*(WT) compared to WT by northern blot hybridization using a probe against the 5’ region of the tRNA precursor. According to transcriptomic data, there are two full-length precursor molecules: One is derived from cotranscription with the gene *slr1256/ureA* (approx. 600 nt). The second, major tRNA^Glu^_UUC_ precursor is 146 nt in length according to transcriptomic data (5’ region: 34 nt, mature tRNA: 73 nt, 3’ region: 39 nt). Signals were detected on both expected sizes (Supp. Fig. S9C) and their intensity was not changed between WT and *rne*(WT), indicating that transcription of the precursors was not affected by RNase E overexpression. However, a processing intermediate accumulated in *rne*(WT) compared to WT a bit below the 100 nt marker band (Supp. Fig. S9C).

**Figure 4:**
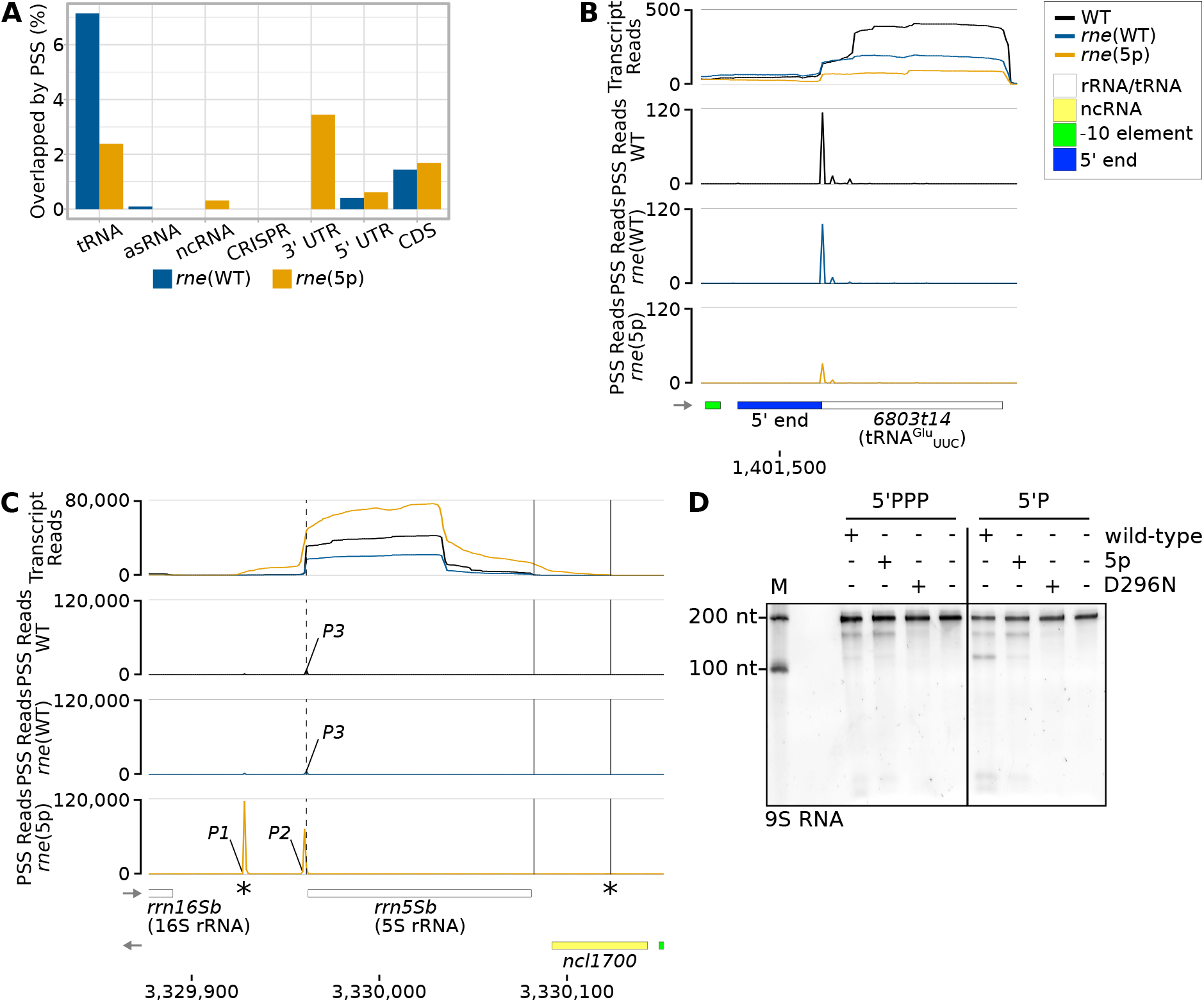
Impact of RNase E on rRNA and tRNA maturation. (**A**) Percentages of different types of transcripts associated with PSS identified in *rne*(WT) and *rne*(5p) relative to the total number of the respective feature type in the annotation. (**B**) RNA-seq data for tRNA^Glu^_UUC_ *(6803t14).* (**C**) RNA-seq data for 5S rRNA *(rrn5Sb).* Dashed vertical lines indicate the 5S rRNA 5’ end detected in WT and *rne*(WT). Solid vertical lines indicate the 3’ ends of 5S rRNA detected in WT and *rne*(WT) (left) or *rne*(5p) (right). Stars (*) indicate transcript boundaries used for *in vitro* cleavage assays (see D). In panels B and C, transcriptome coverage is given on top for the three indicated strains. Cleavage sites P1, P2 and P3 are displayed in the diagrams underneath by black, blue and orange peaks, representing 5’-P (PSS) RNA ends. Transcriptome coverage and PSS represent the average of normalised read counts of the investigated replicates for the indicated strains. For further details, see Supplemental Results S2. (**D**) *In vitro* cleavage assay of a 209 nt 5S rRNA precursor equivalent to the *E. coli 9S* RNA. Transcript boundaries were determined according to RNA-seq transcript coverage (see C). Assays were conducted on transcripts with either 5’-triphosphorylated (5’PPP) or 5’-monophosphorylated (5’P) ends with *Synechocystis* wild-type RNase E (wild-type), or enzymes harbouring the amino acid exchanges T161V (5p) or D296N (D296N). The latter renders RNase E catalytically inactive. One representative urea-PAA gel is shown (n=3). M: marker.

**Table 5:** Processing sites (PSS) in tRNAs differentially regulated between *rne*(WT) and *rne*(5p) with |log_2_FC|>1.5 and adjusted p values <0.05. The corresponding transcriptional units (TU) are indicated according to Kopf et al. (43). The PSS are sorted according to 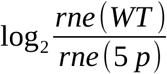.

| Locus tag | TU | tRNA | comment | $\log_2\text{FC}$ | $p_{\text{adj}}$ |
| --- | --- | --- | --- | --- | --- |
| *6803t13_3' UTR* | TU1364 | tRNA<sup>Asp</sup><sub>GUC</sub> | PSS is located 1 nt from 3' end | -2.74 | 8.77E-06 |
| *6803t26* | TU2321 | tRNA<sup>Ala</sup><sub>GGC</sub> | PSS is precisely at 5' end | 1.51 | 0.004 |
| *6803t02* | TU241 | tRNA<sup>Ser</sup><sub>UGA</sub> | PSS is located 1 nt from 3' end | -6.16 | 0.0057 |
| *6803t36* | TU3152 | tRNA<sup>Gly</sup><sub>GCC</sub> | PSS is precisely at 5' end | -2.67 | 0.0147 |
| *6803t14* | TU1442 | tRNA<sup>Glu</sup><sub>UUC</sub> | Figure 4B; this is the major tRNA for all tetrapyrrol biosynthesis, processing is precisely at 5' end | 1.62 | 0.0149 |
| *6803t13_3' UTR* | TU1364 | tRNA<sup>Asp</sup><sub>GUC</sub> | PSS is located 71 nt 3' of tRNA end. Read-through from the tRNA into an asRNA to *slr1740* | -3.98 | 0.0367 |
| *6803t32* | TU2719 | tRNA<sup>Ala</sup><sub>CGC</sub> | PSS is precisely at 5' end | 1.93 | 0.04 |

Several 5’-sensing-dependent PSS mapped to the precursor transcript from which 16S rRNA, the tRNA^Ile^GAU, 23S rRNA and 5S rRNA are matured (Fig. 4C, Supp. Results S2). We observed 5S rRNA precursors accumulating in the 5’ sensing mutant both in urea-PAA gels of total RNA stained with ethidium bromide and by northern blot hybridization (Fig. 1D and E). We performed sequence and structure comparisons between *E. coli* and *Synechocystis* 5S rRNA precursors (Supp. Results S2, Supp. Fig. S10). Based on these and RNA-seq data presented above, RNase E seems to act on a 5S rRNA precursor beginning 34 nt upstream of mature 5S rRNA (PSS *P1*), leading to a precursor which is one nt longer than the mature molecule (*P2* and *P3*). The respective cleavage corresponds to the *E. coli* cleavage site *a* within the 9S RNA, one of the 5S rRNA precursors (Supp. Fig. S10) (62, 63). Also, this analysis indicated an RNase E cleavage event downstream of mature 5S rRNA, reminiscent of the *E. coli* 9S RNA cleavage site *b* (Supp. Fig. S10) (62, 63). Through *in vitro* cleavage assays, we verified the direct action of RNase E on the *Synechocystis* 5S rRNA precursor, which corresponds to the *E. coli* 9S RNA (Fig. 4D). Notably, we also observed 5’-sensing-dependent PSS within mRNAs encoding several ribosomal proteins (*rps21, rps14, rps1a, rps20, rpl27* and *rpl31*) (Supp. Table S10).

### Single examples highlight the role of 5’ sensing in various RNA processing events

There are several studies suggesting strongly that cyanobacterial RNase E autoregulates its own transcript via its 5’ UTR (32, 33, 56, 64), similar to *E. coli* (65). We observed a 5’-sensing-dependent PSS in the 5’ UTR of *rne* (pos. 92,762; Fig. 5A). The accumulation of PSS in *rne*(5p) relative to *rne*(WT) suggests that 5’ sensing plays an important role in further processing of the respective RNA fragment. A recent study mapped RNase E cleavage sites to exactly this position within an A and U rich region by combining 3’ RACE and *in vitro* cleavage assays (56). Furthermore, we mapped RNase-E-dependent cleavage sites a bit downstream using TIER-seq (pos. 92,771; 92,788; 92,797; 92,801) (33). Despite the PSS accumulating in *rne*(5p) compared to *rne*(WT), we did not observe an effect on downstream transcript levels of *rne* and *rnhB* coding sequences (Fig. 5A).

**Figure 5:**
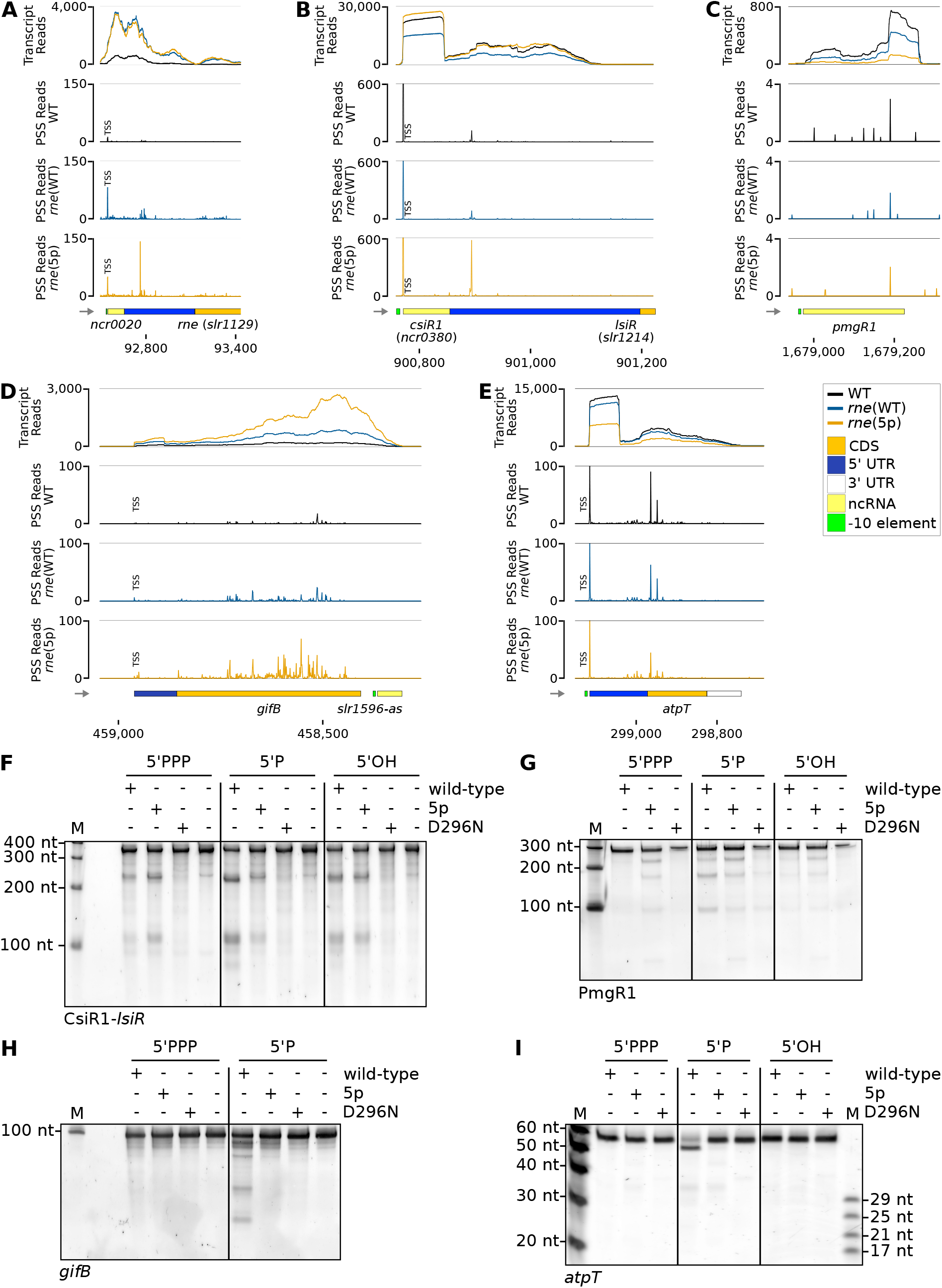
Examples of 5’ sensing dependent cleavage events. RNA-seq data for:(**A**) The 5’ UTR of *rne*. Low read counts surrounding the 5’ end of *rne* are an artefact due to the introduced 3xFLAG sequence in *rne*(WT) and *rne*(5p), which is missing in the sequence to which reads were mapped. (**B**) The CsiR1-*lsiR* transcript. (**C**) The *pmgR1* gene. (**D**) The *gifB* gene including its 5’ UTR and the asRNA located downstream, *slr1596-as.* (**E**) The gene *atpT* including its 5’ and 3’ UTR. Panels (F) to (I): *In vitro* cleavage assays comparing processing of several transcripts presented either 5’-triphosphorylated (5’PPP), 5’-monophosphorylated (5’P) or 5’-hydroxylated (5’OH) by *Synechocystis* wild-type RNase E (wild-type), or enzymes harbouring the amino acid exchanges T161V (5p) or D296N (D296N). The latter renders RNase E catalytically inactive. M: marker. Representative results of 3 to 4 replicates are shown for: (**F**) The 5’ UTR of *lsiR*, including sRNA CsiR1. (**G**) The sRNA PmgR1. (**H**) The 5’ UTR of *gifB.* (**I**) The region around the *atpT* start codon. The substrate consists of the last 12 nt of the 5’ UTR and the first 38 nt of the *atpT* coding sequence. Hence, the two peaks seen in panel (E) are included in the substrate. For panels (A) to (E), transcriptome coverage is given on top for the three indicated strains. Cleavage sites are displayed in diagrams below by black, blue and orange peaks, representing 5’-P (PSS) detected in the three different strains. 5’-PPP (TSS) may be converted to 5’-P RNA ends *in vivo* or during RNA-seq library preparation. Thus, TSS are partially detected in the PSS signal. Positions which were classified as TSS are indicated by “TSS” next to the respective peaks. Transcriptome coverage and cleavage sites (PSS) represent the average of normalised read counts of the investigated replicates. CDS: coding sequences, UTR: untranslated region, ncRNA: non-coding RNA.

The CsiR1-*lsiR* transcript comprises the sRNA CsiR1 (SyR14, ncr0380) and mRNA *lsiR* (*slr1214*), encoding the CheY-like PATAN domain response regulator LsiR. The transcript level is responding to the availability of inorganic carbon, UV-A illumination and ethylene (66–69). LsiR plays a role in the control of negative phototaxis (67–69). Under certain growth conditions, CsiR1 accumulates differentially to the downstream mRNA encoding *lsiR* (43). A 5’-sensing dependent PSS located in the region between CsiR1 and *lsiR* indicates that RNase E might play a role in the differential accumulation of both parts of the transcript (Fig. 5B). Indeed, the 5’ UTR of *lsiR*, including CsiR, was also an *in vitro* target of RNase E and showed a changed processing pattern dependent on its 5’ phosphorylation status (Fig. 5F).

Further potential targets of RNase E included PmgR1, the 5’ UTR of *gifB* and the *atpT* transcript (Fig. 5C, D and E), which substantiates previously published results (33). PmgR1 is an sRNA required for photomixotrophic growth (70). The gene *gifB* encodes the glutamine synthetase inactivating factor IF17. Notably, its 5’ UTR contains a glutamine riboswitch (71). The *atpT* transcript encodes the cyanobacterial ATP synthase inhibitory factor AtpΘ (72). When tested *in vitro*, all three RNA transcripts were cleaved by RNase E in a 5’ phosphorylation dependent manner and cleavage patterns differed between wild-type RNase E and RNase E with the amino acid exchange T161V (Fig. 5G, H, I). For the *atpT* transcript, post-transcriptional regulation was identified as the main level of control (73). The observed 5’P-dependent cleavage occurs at position 298,948 (Supp. Tables S10 and S12), 25 nt downstream of the start codon, effectively inactivating the mRNA.

The genes *gifB, gifA* and *pirA* encode small protein regulators of nitrogen assimilation (74–76); therefore, their identification as RNase E targets adds this RNase to the elements that control this important part of primary metabolism.

## Discussion

### 5’ sensing as an important factor for RNA target recognition for *Synechocystis* RNase E

Systematic studies on the prevalence and relevance of the 5’ sensing function of RNase E have rarely been performed. To our knowledge, the only transcriptome-wide study exploring the impact of 5’ sensing was performed by Clarke et al. (8). They incubated total *E. coli* RNA with the N-terminal half of *E. coli* RNase E harbouring the amino acid exchange T170V *in vitro.* They identified 5’-sensing-dependent RNase E cleavage sites by performing RNA-seq on samples taken before and after the incubation.

Here, engineering of a 5’ sensing deficient RNase E mutant in *Synechocystis* enabled us to investigate the role of 5’ sensing in cyanobacteria *in vivo.* Homozygous clones carrying the mutation leading to T161V were only obtained when the *rne*-*rnhB* operon was expressed from a plasmid vector, but not following integration into a neutral site in the chromosome. Hence, a higher expression level was required, pointing at an important role of 5’ sensing for enzyme function. Moreover, all transcripts selected for *in vitro* analysis showed a changed processing pattern between the wild-type and 5’-sensing-deficient enzyme and depending on the 5’ phosphorylation status. This underlines the role of 5’ sensing for enzyme function and cleavage site positioning.

However, compared to the impact of transient RNase E inactivation (33), only a small number of PSS changed when comparing *rne*(5p) and control strain *rne*(WT). This is in line with findings in *E. coli* that many transcripts can be cleaved by RNase E irrespective of their 5’-phosphorylation status (8). Notably, PSS associated with the maturation of crRNAs from the CRISPR3 array were not affected by the introduced point mutation, albeit responding strongly to RNase E overexpression in both *rne*(WT) and *rne*(5p) compared to WT as well as to transient RNase E inactivation (33). The enhanced levels of these matured crRNA species are consistent with RNase E as the reported single-most important limiting factor for their accumulation (29).

Conversely, the action of RNase E on certain other substrates, such as the *rne* 5’ UTR, was impeded by the mutation present in *rne*(5p). Thus, alternative target recognition determinants also seem to exist for *Synechocystis* RNase E, despite the lack of the long C-terminal half typical for *E. coli* RNase E, which was speculated to be involved in respective recognition mechanisms (5, 15, 24). Notably, protein-protein interaction sites with DEAD-box helicase CrhB and RNase II were mapped to the N-terminal, catalytic domain of RNase E of the cyanobacterium *Anabaena (Nostoc)* sp. PCC 7120 (77, 78). In *E. coli*, the analogous binding site of helicase RhlB is located within the C-terminal domain (31). Hence, despite the compact character of RNase E, many aspects of *E. coli* RNase E seem to be conserved in cyanobacterial RNase E homologues and are fulfilled by regions within the N-terminal half of the enzyme.

### Evidence for plasmid copy number control by RNase E

In our RNA-seq analysis, we observed a general upregulation of plasmid-encoded transcripts in *rne*(WT) and in *rne*(5p) compared to WT. This is most pronounced for pSYSA and pSYSM, which harbour together a total of 232 protein-coding genes (79) and are transcribed in 229 discernible transcriptional units (43). Searching for a common reason, we found an increased copy number of these two plasmids relative to the chromosome, with an up to ~2.4 higher copy number for pSYSM and a ~3.8 fold higher copy number for pSYSA (in both cases comparing *rne*(WT) to WT and pVZΔKm^R^). Somewhat lower, but also clearly increased copy numbers, were observed when comparing *rne*(5p) to WT and pVZΔKm^R^. We conclude that overexpression of RNase E and RNase HII from the introduced *rne-rnhB* operon leads to increased pSYSA and pSYSM copy numbers. *E. coli* homologues of both enzymes play a decisive role in the copy number regulation of ColE1 and ColE2-type plasmids by processing pairs of overlapping non-coding RNAs, of which one can then prime replication (80–82). Indeed, we identified a similar pair of abundant overlapping transcripts on pSYSA: *ssr7036* mRNA and asRNA1 (83). Both transcripts were verified as substrates for RNase E and their manipulation led to divergent pSYSA copy numbers (83). There is a growing interest in repurposing the endogenous plasmids of cyanobacteria for their efficient metabolic engineering. Hence, the here gained insight in their copy number control mechanisms will be useful.

### Inactivating 5’ sensing *in vivo* enables to identify RNase E targets and to elucidate roles of RNase E in cyanobacteria

Analysing the impact of decreased 5’ sensing, we identified previously unknown targets of and processing events dependent on RNase E in *Synechocystis.* Changes in rRNA and tRNA maturation in the 5’ sensing RNase E mutant demonstrate that these functions of RNase E known from analyses in *E. coli* (5, 7, 9, 84) are conserved in cyanobacteria. Furthermore, mapping of 5’-sensing-dependent PSS to CDS of ribosomal proteins and the general impact of *rne-rnhB* overexpression on ribosomal protein transcript levels observed in GSEA hint towards *Synechocystis* RNase E directly regulating transcript levels of ribosomal proteins. This is consistent with *E. coli* RNase E acting on *rpsO* and *rpsT* mRNAs, which encode the 30S ribosomal proteins S15 and S20 (5, 85). Also, in *Arabidopsis thaliana* chloroplasts, RNase E was found to have an important function in processing of ribosomal protein mRNAs (86).

Recently, Mohanty and Kushner (9) found that *E. coli* RNase E plays an important role in the processing of the *alaW alaX* operon. Interestingly, this processing was not abolished in a mutant strain encoding thermosensitive RNase E. Since transcriptome-wide studies identifying RNase-E-dependent processing sites commonly use thermosensitive strains, this implies that these miss a subset of RNase-E-dependent processing events (8, 14, 33, 87). Employing a 5’-sensing-deficient RNase E mutant strain might help to provide a more complete picture of processing sites.

Neither decreased 5’ sensing nor RNase E overexpression did affect PSS abundance or PSS/TSS ratios at TSS positions. This is consistent with our prior finding that RNase E does not seem to be rate-limiting for 5’-end dependent RNA degradation and is further hinting at a central role of RNase J in general RNA degradation, or a redundant action of RNase E and RNase J therein (33).

The majority of RNA features most strongly affected by decreased 5’ sensing are non-coding, potentially regulatory elements, e.g. asRNAs, ncRNAs or 5’ UTRs (Tables 2 and 4). These might be directly regulated by RNase E, consistent with known functions of *Synechocystis* RNase E (26, 28). Further examples for the action of RNase E on ncRNAs are the processing of the CsiR1-*lsiR* actuaton, PmgR1 and the 5’ UTRs of both *gifB* and *rne*. Although the exact mechanisms and physiological impacts of these cleavage events remain to be elucidated, these findings hint strongly at a general regulatory role of RNase E in cyanobacteria. The analysed data set maps many further interesting RNase-E-dependent cleavage sites, which await further characterization, e.g. a site located within the *apcABCZ* operon upstream of the sRNA ApcZ (88) and within the *grpE* gene (Supp. Results S3, Supp. Fig. S11).

Moreover, we show that RNase E is involved in the maturation of several tRNAs *in vivo.* The involvement of RNase E in the initial processing of tRNA precursors was reported in *E. coli*, where RNase E activity provides the substrates for further maturation by RNase P and the 3’ exoribonucleases (89). The most striking example for RNase E involvement in the maturation of tRNA precursors is the one-step endonucleolytic cleavage immediately after the CCA determinant yielding mature tRNAs for three different proline tRNA species in *E. coli (90, 91).* Here, we present evidence for the involvement of RNase E in the accurate 5’ maturation of the glutamyl-tRNA tRNA^Glu^_UUC_ by RNase E. This particular tRNA is of greater interest because it has a twofold role in *Synechocystis:* In addition to its function in translation, it activates the amino acid glutamate as the precursor of tetrapyrrole synthesis. Hence, all cytochromes, tetrapyrroles and, especially heme and chlorophylls derive from glutamate activated via binding to this tRNA (60, 92, 93). This pathway is present in cyanobacteria, plants, many bacteria and Archaea. In *Synechocystis*, this pathway was demonstrated as the sole biosynthetic pathway for tetrapyrrole biogenesis and there is only one corresponding glutamyl-tRNA gene present in the genome (61, 94, 95). It is very well established that mature tRNA 5’ ends are generated by RNase P (96). So, why would RNase E have evolved in certain cyanobacteria to participate in the maturation of this tRNA? Previously, a deviant consensus was described for the maturation of the tRNA^Glu^_UUC_ 5’ end generated by RNase P in cyanobacteria and chloroplasts. This tRNA differs in cyanobacteria and chloroplasts from the eubacterial tRNA consensus by the presence of a unique A53:U61 instead of the otherwise canonical G53:C61 base pair at the distal end of the T stem (60, 61). The participation of the T stem in substrate binding to the RNase P ribozyme has been established for RNase P RNA from various structural classes (97, 98). It was demonstrated that this difference makes the cyanobacterial and chloroplast tRNAs less suitable substrates for RNase P from these organisms and that the experimental replacement of A53:U61 by G53:C61 improved product formation (99, 100).

Our findings suggest that the 5’ maturation of tRNA^Glu^_UUC_ in *Synechocystis* is supported by RNase E. Plant chloroplasts harbour a close homolog to the cyanobacterial glutamyl-tRNA as well as RNase E (30). Therefore, the investigation of this relation and its physiological relevance beyond *Synechocystis* or cyanobacteria is an interesting topic for future research. Generally, the RNase E from *Synechocystis* is a great model to study RNA maturation and degradation in bacteria as it can be produced efficiently as an enzymatically active recombinant protein and appears to share substantial substrate preferences with its enterobacterial homologues despite its more compact nature. Exploring the phylogenetic diversity of different RNase E homologues and inactivating specifically 5’ sensing will help to elucidate how this multi-faceted enzyme recognises its target molecules.

## Data availability

FastQ files from RNA sequencing have been deposited with the Gene Expression Omnibus (GEO) Data bank under accession number GSE184824. Galaxy workflows used for raw read processing and several downstream analysis steps are available online: https://usegalaxy.eu/u/ute-hoffmann/w/5sens-seq-preprocessing, https://usegalaxy.eu/u/ute-hoffmann/w/5sensing-pss-tss, https://usegalaxy.eu/u/ute-hoffmann/w/5sensing-transcript, https://usegalaxy.eu/u/ute-hoffmann/w/5sensing-transcript-coverage

All further code used for data processing and analysis are available online in the GitHub and Zenodo repositories (https://github.com/ute-hoffmann/5sensing_Synechocystis, https://doi.org/10.5281/zenodo.7509658).

## Supplementary data

Supplementary Data are available online.

- 5sensing_paper_supplement.pdf
- SupplementaryTables_S5-S14.ods

## Acknowledgements

We thank J. Behler, T. Wallner, P. Schwenk, M. Treppner and N. Schuergers for fruitful discussions. We thank W. Bigott for technical assistance. The authors acknowledge the support of the Freiburg Galaxy Team: Björn Grüning, Bioinformatics, University of Freiburg (Germany) funded by the Collaborative Research Centre 992 Medical Epigenetics (Deutsche Forschungsgemeinschaft [SFB 992/1 2012]) and Bundesministerium für Bildung und Forschung BMBF grant 031 A538A de.NBI-RBC.

