## Supplemental Information for "The role of the 5’ sensing function of ribonuclease E in cyanobacteria"

### The targetome of a cyanobacterial RNase E deficient in 5' sensing

#### Supplemental Information

**Supplemental Methods**

**Supplemental Results**

**Supplemental Tables**

**Supplemental Figures**

**Supplemental References**

#### Supplemental Methods

##### LocARNA alignment

Sequence-structure alignments were performed using the LocARNA (v1.9.1, linking Vienna RNA package v2.3.2) webserver with default parameters (Raden et al. 2018; Will et al. 2007; 2012).

##### Quantitative PCR

DNA of three independent cultures of WT and *rne*(5p) as well as four independent cultures of pVZΔKm$^{R}$ and *rne*(WT) were used for quantifying copy numbers of *Synechocystis* stable plasmids pSYSA, pSYSM, pSYSX, pSYSG and the introduced conjugative, self-replicating plasmid pVZ321 relative to chromosomal DNA using qPCR and primers qPCR1 to qPCR12 described in Supplementary Table 1. Primer specificity was analysed using Primer Blast (Ye et al. 2012).
qPCR reactions were set up according to the QuantiFAST-SYBR-Green-PCR manual (QIAGEN, Netherlands). Optimal primer concentrations and the occurrence of non-specific PCR products were tested by melting curve analyses and verified by agarose gel electrophoresis. For plasmid quantification, 1 µM of each primer was used and approximately 5 ng of total DNA as template in 10 µl reactions. Reactions were set up in 384 FastGene plates (Nippon Genetics Europe GmbH, Germany) and measured on a CFX384 Touch real-Time PCR Detection System (Bio-Rad Laboratories, USA). The cycling conditions were set to 5 min at 95°C, followed by 40 cycles of 10 s denaturation at 95°C and 30 s annealing/extension at 60°C. A melting curve analysis was performed for each qPCR run to identify secondary products. For calibration of the quantification of starting quantities, a dilution series encompassing six dilution steps between 0.1 ng and 100 ng was assayed to analyse amplification efficiency for each primer pair. Biological replicates were measured in technical triplicates. After calculation of starting quantities, plasmid relative to chromosomal quantity was calculated.

#### Supplemental Results

##### S1. Comparison of 5'-sensing-dependent PSS to RNase-E-dependent PSS identified by TIER-seq

Recently, we mapped RNase-E-dependent PSS on a transcriptome-wide level using the TIER-seq approach (Hoffmann et al. 2021). To do so, we compared PSS in *rne*(WT) and in a strain harbouring temperature-sensitive RNase E (*rne*(Ts)) after a transient incubation at 39°C, which inactivates RNase E in *rne*(Ts). PSS which are less abundant after heat inactivation of RNase E are bona fide cleavage sites of the enzyme. PSS with more counts after RNase E heat inactivation indicate RNA fragments which could accumulate because they are not degraded due to the lowered ribonuclease activity. We re-analysed the combined set of all PSS identified here with the original TIER-seq PSS. Of this common set, 731 PSS accumulated in *rne*(WT) and 717 in *rne*(Ts) after the heat treatment. The majority of PSS accumulating in *rne*(WT) compared to *rne*(5p) here overlap with PSS which accumulated also previously in *rne*(WT) in the TIER-seq data set (62 sites, 59.0%, Supp. Fig. 5). Similarly, PSS with higher counts in *rne*(5p) share the highest overlap of 63 sites with PSS accumulating in *rne*(Ts) after the heat shock (35.4%). Such RNase-E-dependent PSS identified in both data sets are highlighted with an asterisk (*) in Table 3.

##### S2. Comparison of rRNA processing in WT, *rne*(WT) and *rne*(5p)

In several species, RNase E was shown to be involved in rRNA maturation (Ghora and Apirion 1979; Li, Pandit, and Deutscher 1999; Taverniti et al. 2011; Klein and Evguenieva-Hackenberg 2002). The action of paralogous *E. coli* RNase E and RNase G on the 5' end of 16S rRNA precursors and the respective cleavages' dependence on 5' sensing is well studied (Li, Pandit, and Deutscher 1999; Garrey and Mackie 2011). Here, we observed a PSS corresponding to the mature 16S rRNA, which did not respond to 5' sensing inactivation, but no further PSS within the 5' elongated segment of 16S rRNA precursors. However, we observed a PSS downstream of mature 16S rRNA to accumulate strongly in *rne*(5p) compared to *rne*(WT) and WT (genomic pos. 2,452,183 and 3,326,545). This points towards a role of 5' sensing in the release of the downstream located tRNA, 23S rRNA and 5S rRNA from the precursor transcript.

The importance of RNase E in 5S rRNA maturation in *E. coli* is well established and the dependence of the respective cleavage events on 5' sensing or secondary structures was intensively studied (Ghora and Apirion 1979; 1978; Cormack and Mackie 1992; Kaberdin et al. 2000; Garrey et al. 2009; Garrey and Mackie 2011; Roy et al. 1983). In *E. coli*, the precursor transcript 9S RNA results from RNase III cleavage (Bram, Young, and Steitz 1980). This precursor is further processed by RNase E to yield pre-5S rRNA. Subsequently, RNase AM and RNase T process pre-5S rRNA to mature 5S rRNA (Jain 2020; Li and Deutscher 1995). Inactivation of RNase E in a temperature-sensitive mutant strain and a strain with decreased 5' sensing led to accumulation of 5S rRNA precursors (Ghora and Apirion 1979; 1978; Garrey et al. 2009; Garrey and Mackie 2011). Here, we observed a similar phenomenon in *Synechocystis*. Compared to WT and *rne*(WT), two 5S rRNA precursors accumulated in *rne*(5p) (Fig. 1F). Based on our sequencing data, we identified three PSS surrounding the annotated 5S rRNA, called *P1*, *P2* and *P3* (Fig. 4C). The PSS accumulating most strongly in *rne*(WT) and WT is located one nt upstream (*P3*, genomic pos. 3,329,961 for *rrn5Sb*, indicated by a vertical dashed line in Figure 4C) of the annotated 5S rRNA gene. In *rne*(5p), two PSS accumulated one nt (*P2*, genomic pos. 3,329,960) and 34 nt (*P1*, genomic pos. 3,329,928) upstream of the WT-PSS *P3*. Furthermore, we noticed that 5S-rRNA-associated transcript reads ended 45 nt downstream of mature 5S rRNA in *rne*(5p) (genomic pos. 3,330,125), but not in WT or *rne*(WT) (most downstream transcript reads associated with 5S rRNA are indicated by vertical solid lines in Fig. 4C). In WT and *rne*(WT), the majority of transcript reads ended close to the 5S rRNA 3' end. To compare these intermediates to *E. coli* 5S rRNA, we performed a sequence-structure alignment of *E. coli* 9S RNA and *Synechocystis* sequences reaching from the 3' end of 23S rRNA to the 3' end detected in *rne*(5p) (Supp. Methods, Supp. Fig. 9). The obtained structure was in good concordance with the findings of Cormack and Mackie (Cormack and Mackie 1992) for *E. coli* 9S RNA. Based on this comparison, we propose mature *Synechocystis* 5S rRNA to start at the mapped WT-PSS (genomic pos. 3,329,961 for *rrn5Sb*). Since *P1* is not located within a single-stranded RNA segment, we assume this PSS not to be created by RNase E. However, accumulation of this PSS in *rne*(5p) compared to WT and *rne*(WT) indicates that further processing to both *P2* and *P3* is dependent on RNase E 5' sensing. Also, comparison to the processing events in *E. coli* indicated that a PSS dependent on RNase E 5' sensing exists at a similar position as the cleavage position *b* within the *E. coli* 9S RNA. Hence, RNase E 5' sensing seems to be important for removal of the terminator hairpin during 5S rRNA maturation.

**S3. Two further examples of 5’ sensing dependent RNA processing**

There are many further interesting cases of 5’ sensing and RNase-E-dependent PSS. One of these is a PSS upstream of ApcZ in the *apcABCZ* operon (Supp. Fig. 10A). ApcZ is an sRNA regulating orange carotenoid protein (OCP) levels and it is partially overlapping with *apcC* (Zhan et al. 2021). It can be transcribed from its own promoter or as part of the *apcABCZ* transcript (Zhan et al. 2021). By northern blotting, transcript turnover products encompassing ApcZ and different parts of the remaining tetracistronic transcript were identified (Zhan et al. 2021). Here, we observed a 5'-sensing-dependent PSS within the coding sequence of *apcB*, probably leading to a transcript encompassing the intergenic region between *apcB* and *apcC*, the *apcC* mRNA and the sRNA ApcZ (Supp. Fig. 10B). The RNase-E-dependent PSS which maps to the *apcB* gene might lead to a functional RNA fragment derived from the intergenic region between *apcB* and *apcC*, reminiscent of the sRNA sponge SroC in *E. coli* (Miyakoshi, Chao, and Vogel 2015). SroC is derived from the 3'

UTR of mRNA *gltI*, which corresponds to the intergenic region between *gltI* and *gltJ* within the *gltIJKL* operon (Miyakoshi, Chao, and Vogel 2015). Maturation of SroC is dependent on RNase E 5' sensing (Miyakoshi, Chao, and Vogel 2015). Furthermore, its regulatory role on the globally acting sRNA GcvB is mediated by RNase E and not dependent on 5' sensing (Miyakoshi, Chao, and Vogel 2015). Alternatively, the PSS might yield monocistronic *apcC* mRNA or initiate the maturation of 3' end derived sRNA ApcZ.

Another interesting PSS is located upstream of an RNA fragment derived from the 3’ end of *grpE*, which is encoding the GrpE nucleotide exchange factor (Supp. Fig. 10B) (Barthel, Rupprecht, and Schneider 2011). *Synechocystis* GrpE can restore growth in an *E. coli grpE* deletion strain at 42°C, indicating that it can act as co-chaperone for the DnaK heat shock protein (Barthel, Rupprecht, and Schneider 2011). Compared to the rest of the coding sequence of *grpE*, the RNA fragment downstream of the PSS is accumulating strongly in *rne*(5p). In *rne*(5p), the mapped 5' end of this RNA fragment is shifted 3 nt upstream compared to the starts identified in WT and *rne*(WT) (pos. 2,589,108 to pos. 2,589,111). The RNA fragment spans a 70 amino acids long open reading frame (ORF) in frame with the *grpE* ORF and corresponding to the C-terminal β-sheet domain (Barthel, Rupprecht, and Schneider 2011). This β-sheet is not necessary for temperature-dependent GrpE dimerization in *Synechocystis* (Barthel, Rupprecht, and Schneider 2011). Co-crystallization of *E. coli* GrpE and the ATPase domain of DnaK showed that the GrpE β-sheet is interacting with the nucleotide-binding cleft of DnaK (Harrison et al. 1997).

#### Supplemental Tables

**Table S1.** Oligonucleotides used in this study The T7 promoter sequences present in some of the primers are underlined.

| Primer name | Primer sequence | Comment |
|---|---|---|
| C01 | TATCCATGGGCCCAAAACAAATTGTCATTGC | Cloning of pCDF RNase E expression vector |
| C02 | GCGTCGACTTAGTGATGGTGATGGTGATG | Cloning of pCDF RNase E expression vector |
| M01 | GACGGTAATTAATGTTAACTCTGGTTCCTTTAC | Amino acid exchange D296N |
| M02 | AAAGCTTCCGTCGGCTCA | Amino acid exchange D296N |
| M03 | GCTAGTGCGGGTGGAAGCGGAAGATGTGCC | Amino acid exchange T161V |
| M04 | AGTCCCATGCCCGGTGGC | Amino acid exchange T161V |
| T01 | TAATACGACTCACTATAGGGCTTCTGGGTTTCTTCTCAGCAA | Amplification of DNA template for *in vitro* transcript of 9S RNA |
| T02 | AAAAGCGAAACCCTCCAACTAC | Amplification of DNA template for *in vitro* transcript of 9S RNA |
| T03 | TAATACGACTCACTATAGGGAGAAAGTAAATCGTTCATCTTCTCTTTTTG | Amplification of DNA template for *in vitro* transcript of *gifB* UTR (gifB-UTR-T7_fw) (Klähn et al. 2018) |
| T04 | AGTGCGCTCCTAATTGTTATTGAAGG | Amplification of DNA template for *in vitro* transcript of gifB UTR (gifB-UTR_rev) (Klähn et al. 2018) |
| T05 | TAATACGACTCACTATAGGGATGGAGCCGAAGGGCTTCACTGGC | Amplification of DNA template for *in vitro* transcript of PmgR1 |
| T06 | TTCCAAACAACACCCAGAACC | Amplification of DNA template for *in vitro* transcript of PmgR1 |
| T07 | TAATACGACTCACTATAGGGAGTCAGTTCCAATCTGAAC | Amplification of DNA template for *in vitro* transcript of first 110 nt of *psbA2* |
| T08 | CCCACTGACAAAACT | Amplification of DNA template for *in vitro* transcript of first 110 nt of *psbA2* |
| T09 | TAATACGACTCACTATAggGTGCTGGTGAGCGACGGGC | Amplification of DNA template for *in vitro* transcript of CsiR1 and 5' UTR of *lsiR* |
| T10 | GGGATAAGGAGACCTAGGGCCGAG | Amplification of DNA template for *in vitro* transcript of CsiR1 and 5' UTR of *lsiR* |
| T11 | TATTGCATAAGGTCCTGTAGCATCTCAAGTAATTTCATCATAACCTCCGTCCCTATAGTGAGTCGTATTA | Amplification of DNA template for *in vitro* transcript of region around *atpT* start codon |
| T12 | TAATACGACTCACTATAGGG | Amplification of DNA template for *in vitro* transcript of region around *atpT* start codon |

| | | |
|---|---|---|
| qPCR1 | TTTTGCTTCCGTGGCAATG | Detection of chromosomal DNA using qPCR |
| qPCR2 | AGGCGAATGACGGAAACTTG | Detection of chromosomal DNA using qPCR |
| qPCR3 | ATTCGCGCTCGTTGTTCTTC | Detection of pVZ DNA using qPCR |
| qPCR4 | TACAAAGTAGGGTCGGGATTGC | Detection of pVZ DNA using qPCR |
| qPCR5 | TATGTCTTTCGGGGCAAACG | Detection of pSYSA DNA using qPCR |
| qPCR6 | GCGGAAAGAATGGGTTGATACC | Detection of pSYSA DNA using qPCR |
| qPCR7 | AATGGCACCTGGTCAAATCC | Detection of pSYSM DNA using qPCR |
| qPCR8 | ACCCAAGCCAAATGCAATCG | Detection of pSYSM DNA using qPCR |
| qPCR9 | TTTGTCATCCGGGTGAATGC | Detection of pSYSX DNA using qPCR |
| qPCR10 | AAGCCGCCAATGATGTGATC | Detection of pSYSX DNA using qPCR |
| qPCR11 | AGCGACCAAACTGCAATAGC | Detection of pSYSG DNA using qPCR |
| qPCR12 | ACATGGCCGTGTTCATCAAC | Detection of pSYSG DNA using qPCR |
| S01 | TGGAAGCTAAGGCTACAGCG | Southern blot hybridization to test homozygosity of *rne*(WT) and *rne*(5p) |
| S02 | AGGGGTTACAGTCAGCTTGC | Southern blot hybridization to test homozygosity of *rne*(WT) and *rne*(5p) |
| S03 | CCCTAGTCGGTTTTGGCTCT | Amplification of DNA templates for *in vitro* transcripts for Southern blot hybridization |
| S04 | <u>TAATACGACTCACTATAGGGG</u>GTATGTCAGCCGATGGAGT | Amplification of DNA templates for *in vitro* transcripts for Southern blot hybridization |
| S05 | CTTTAGGTGGGCGTTGACCT | Amplification of DNA templates for *in vitro* transcripts for Southern blot hybridization |
| S06 | <u>TAATACGACTCACTATAGGG</u>TAATAGTAATGACAGGCAG | Amplification of DNA templates for *in vitro* transcripts for Southern blot hybridization |
| S07 | TTCCCCTAGCTTCGGCTTTG | Amplification of DNA templates for *in vitro* transcripts for Southern blot hybridization |
| S08 | <u>TAATACGACTCACTATAGGG</u>AACGGTCCGAGAATCAGCAG | Amplification of DNA templates for *in vitro* transcripts for Southern blot hybridization |

| | | |
|---|---|---|
| N01 | TGGTGGTGTCCACGGGCA | Amplification of DNA templates for *in vitro* transcripts for Northern blot hybridization |
| N02 | <u>TAATACGACTCACTATAGGG</u>GACGGGAAAGATTCACTCCCC | Amplification of DNA templates for *in vitro* transcripts for Northern blot hybridization |
| N03 | GTCTGAAATAACGAACTGTTATCGAGACTGCCTA<u>CCCTATAGTGAGTCGTATTA</u> | *In vitro* transcription of probe complementary to tRNA 5’ end |
| N04 | <u>TAATACGACTCACTATAGGG</u> | *In vitro* transcription of probe complementary to tRNA 5’ end |
| 5S-rRNA-oligo | GCATCGGACTATTGTGCCGTG | Oligonucleotide to detect 5S rRNA |

**Table S2.** Strains used in this study.

| Name | Description |
|---|---|
| pVZΔ$Km^R$ (Hoffmann et al. 2021) | *slr1128*::*kanR 3xFLAG-*$rne^{+}$ pVZ321Δ(9220-1203) |
| 3xFLAG-*rne* (Hoffmann et al. 2021) | *slr1128*::*kanR 3xFLAG-*$rne^{+}$ |
| *rne*(WT) (Hoffmann et al. 2021) | Δ(*rne-rnhB*) *rnep*::*kanR* pVZ321Δ(9,220-1,203) (9,219)::(*rnep-3xFLAG-*$rne^{+}$*-rnhB-rnhBt*) |
| *rne*(5p) | Δ(*rne-rnhB*) *rnep*::*kanR* pVZ321Δ(9,220-1,203) (9,219)::(*rnep-3xFLAG-rne*(5p)*-rnhB-rnhBt*) |

**Table S3.** Gene set enrichment analysis (GSEA) using KEGG terms, comparing *rne*(WT) and WT. Higher values are enriched in *rne*(WT).

| ID | Description | Set size | Enrichment score | p.adj |
|---|---|---|---|---|
| syn03010 | Ribosome | 53 | 0.62 | 7.5E-04 |
| syn00910 | Nitrogen metabolism | 18 | -0.74 | 2.5E-03 |
| syn00920 | Sulfur metabolism | 14 | 0.76 | 3.0E-02 |
| syn00190 | Oxidative phosphorylation | 49 | -0.51 | 3.0E-02 |

**Table S4.** Gene set enrichment analysis (GSEA) using KEGG terms, comparing *rne*(WT) and *rne*(5p). Higher values are enriched in *rne*(WT).

| ID | Description | Set size | Enrichment score | p.adj |
|---|---|---|---|---|
| syn03010 | Ribosome | 53 | -0.63 | $8.1 * 10^{-4}$ |

**Overview further supplementary Tables**

Further Supplementary Tables are available in a separate file (SupplementaryTables_S5-S14.ods). They encompass:

**Table S5.** DESeq2 analysis on level of RNA features comparing WT and *rne*(WT). RNA features are sorted according to increasing adjusted p values.

**Table S6.** DESeq2 analysis on level of RNA features comparing *rne*(WT) and *rne*(5p). RNA features are sorted according to increasing adjusted p values.

**Table S7.** DESeq2 analysis on level of transcriptional units comparing WT and *rne*(WT). Transcriptional units are sorted according to increasing adjusted p values.

**Table S8.** DESeq2 analysis on level of transcriptional units comparing *rne*(WT) and *rne*(5p). Transcriptional units are sorted according to increasing adjusted p values.

**Table S9.** DESeq2 analysis on level of PSS comparing WT and *rne*(WT). PSS are sorted according to increasing adjusted p values.

**Table S10.** DESeq2 analysis on level of PSS comparing *rne*(WT) and *rne*(5p). PSS are sorted according to increasing adjusted p values.

**Table S11.** DESeq2 analysis on level of PSS comparing WT and *rne*(WT), including multi-mapping reads. PSS are sorted according to increasing adjusted p values.

**Table S12.** DESeq2 analysis on level of PSS comparing *rne*(WT) and *rne*(5p), including multi-mapping reads. PSS are sorted according to increasing adjusted p values.

**Table S13.** Replicon gene set enrichment analysis comparing WT and *rne*(WT). Higher values are enriched in *rne*(WT).

**Table S14.** Replicon gene set enrichment analysis comparing *rne*(WT) and *rne*(5p). Higher values are enriched in *rne*(WT).

#### SUPPLEMENTAL FIGURES

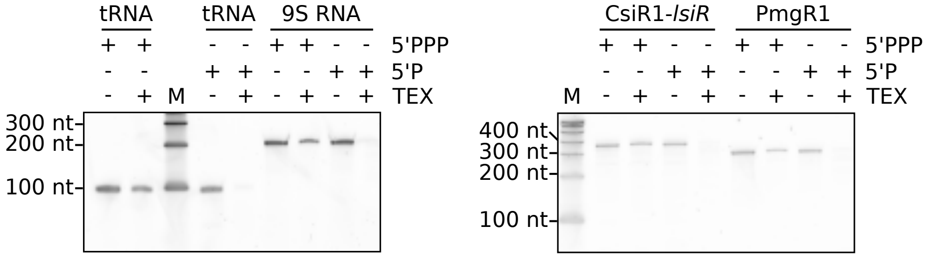


**Supplementary Figure S1.** Confirmation of phosphorylation status of several *in vitro* transcripts using TEX. The 5'-triphosphorylated (5'PPP) transcripts were generated by *in vitro* transcription. The 5'PPP ends were converted to 5'-monophosphorylated (5'P) by incubating transcripts with RppH. M: marker

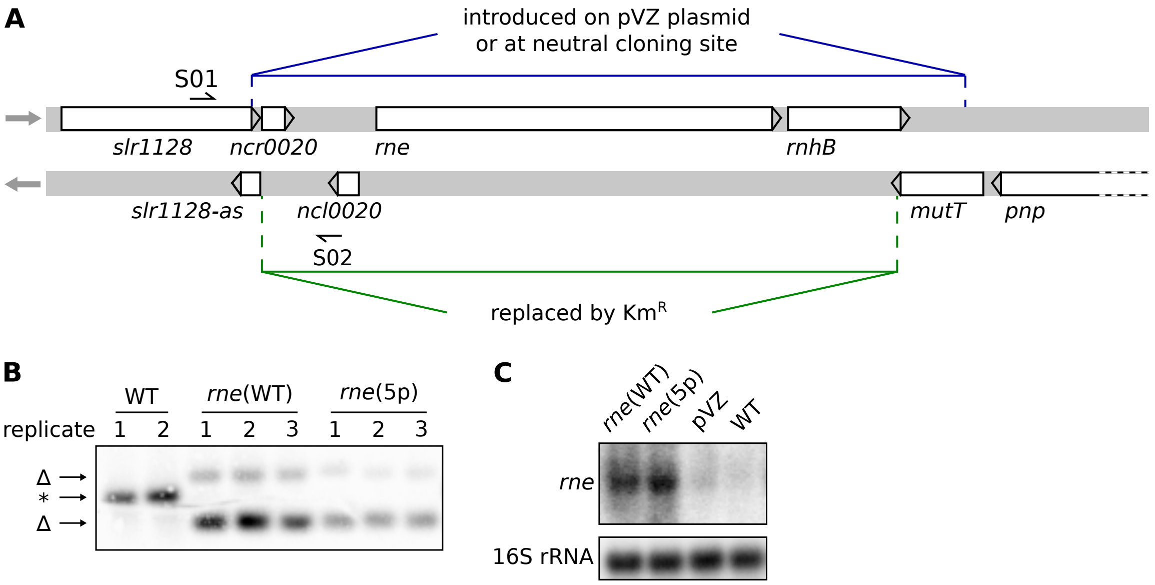


**Supplementary Figure S2.** Construction of *rne-rnhB* mutant strains. (**A**) Overview of the *rne-rnhB* genomic locus. (**B**) Southern blot to test homozygosity of mutant strains. The membrane was hybridised with a probe generated using primers S01 and S02. If the locus was not disrupted by insertion of the kanamycin resistance cassette, only one band occurs (WT, indicated by asterisk *). After insertion, the locus is disrupted and two bands occur (indicated by $\Delta$). (**C**) Representative northern blot analysis using a probe for the coding sequence of *rne* (n=3).

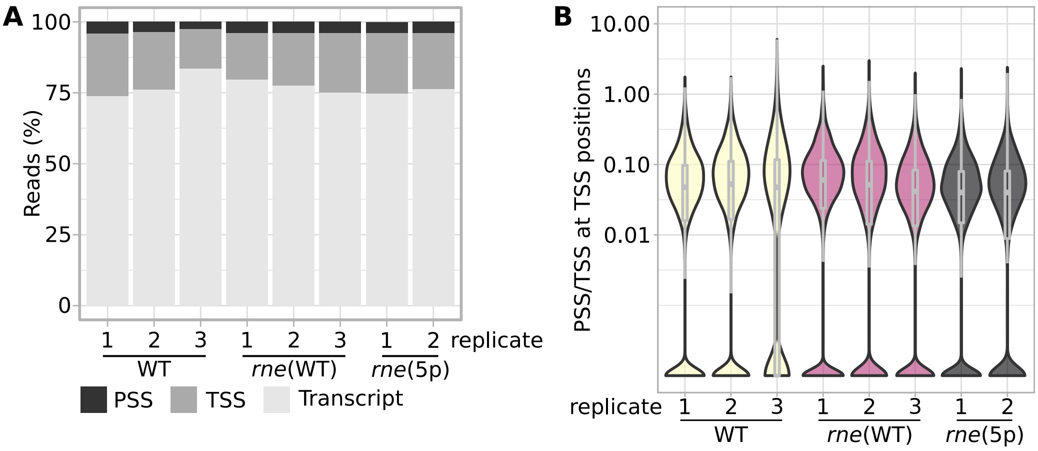


**Supplementary Figure S3.** Ratio of PSS and TSS reads in different samples. (**A**) Shares of reads corresponding to PSS, TSS or unspecified transcript positions in the 8 samples. (**B**) Ratio of PSS to TSS counts at TSS positions in the 8 samples.

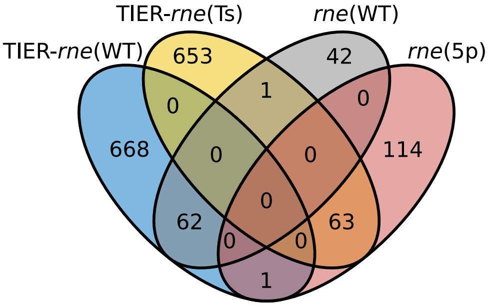


**Supplementary Figure S4.** Comparison of the RNase-E-dependent PSS identified here and in a previous TIER-seq analysis (Hoffmann et al. 2021). The sets of PSS accumulating in the data set presented here in *rne*(WT) or *rne*(5p) are labelled with *rne*(WT) and *rne*(5p), respectively. With TIER-seq, a temperature-sensitive RNase E mutant strain (*rne*(Ts)) was compared to the control strain *rne*(WT) after 1 h incubation at the non-permissive temperature of 39°C (Hoffmann et al. 2021). PSS identified by comparison of both strains after 1 h are labelled TIER-*rne*(WT) and TIER-*rne*(Ts) for PSS accumulating in *rne*(WT) and *rne*(Ts), respectively.

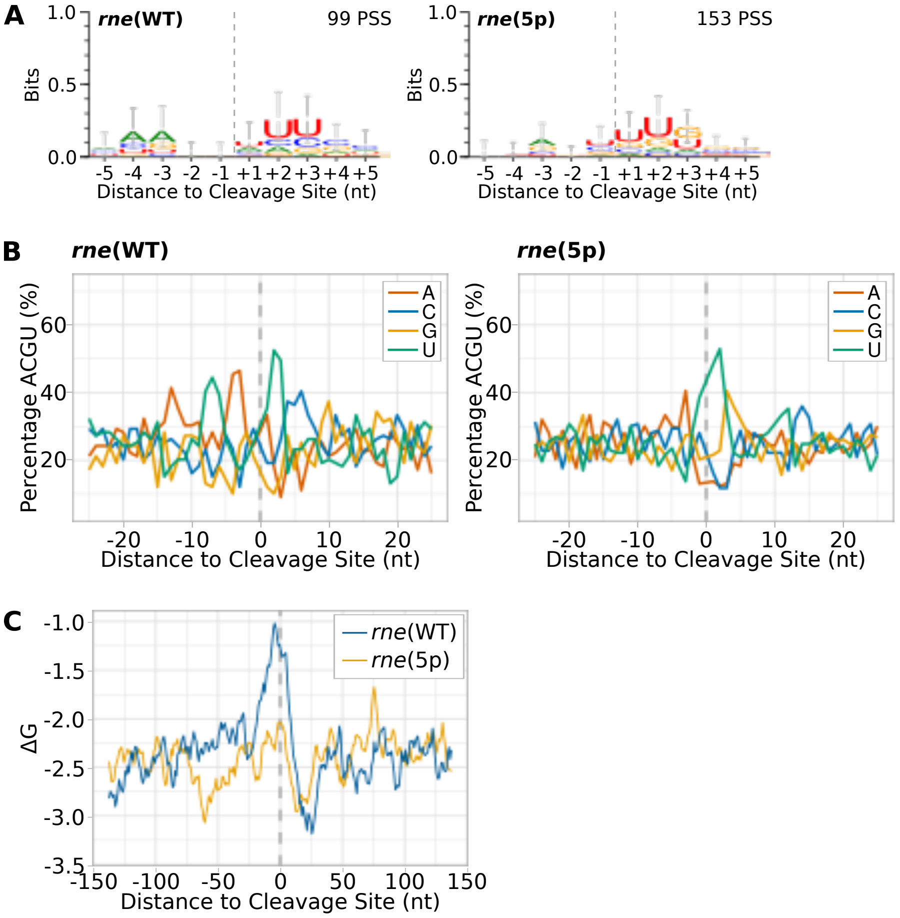


**Supplementary Figure S5.** Inference of sequence signature and folding potential around cleavage sites. (**A**) Sequence logos for PSS higher in either *rne*(WT) (left) or *rne*(5p) (right). Sequences were aligned according to the detected cleavage site. Error bars were calculated by the WebLogo tool and represent an approximate Bayesian 95% confidence interval. (**B**) Nucleotide content surrounding PSS higher in *rne*(WT) or *rne*(5p). (**C**) Minimal folding energy ($\Delta$G) in regions 150 nt up- and downstream of PSS with higher counts in *rne*(WT) or *rne*(5p). Minimal folding energy was calculated at each nucleotide position using a sliding window encompassing 25 nt.

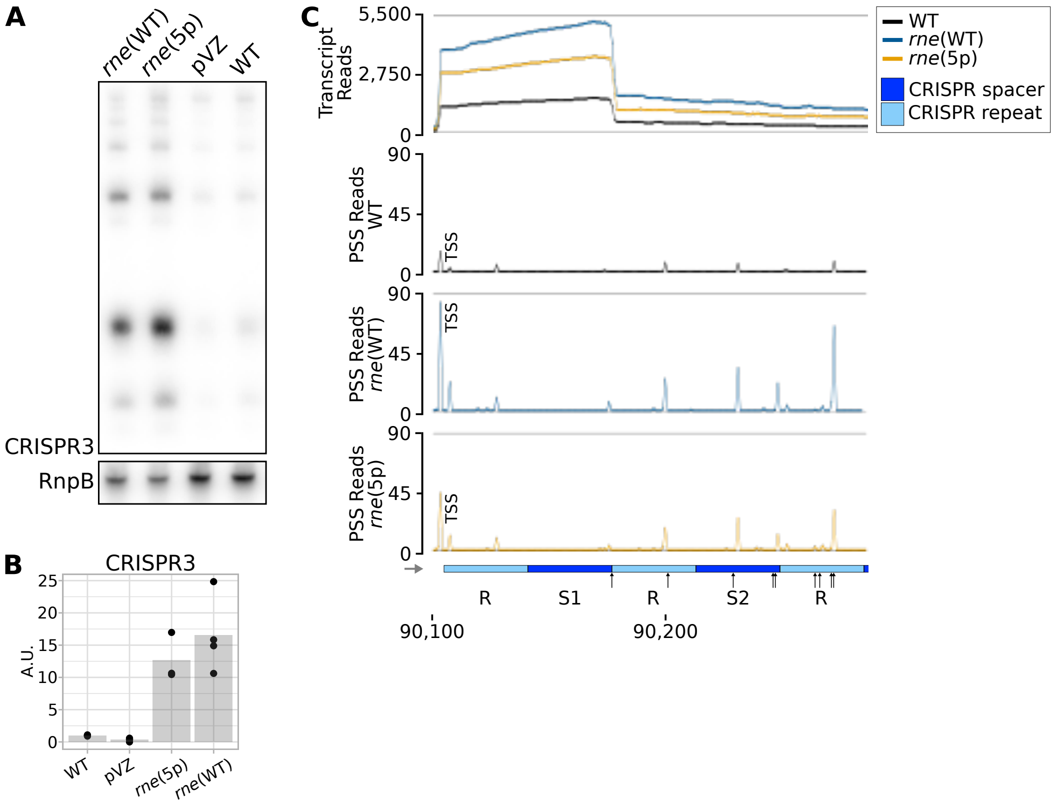


**Supplementary Figure S6.** (**A**) Representative northern blot analysis of CRISPR3 accumulation (n=3). (**B**) Quantification of CRISPR3 hybridization signal. (**C**) RNA-seq data for spacers 1 and 2 of the CRISPR3 array. The black arrows point at positions at which cleavage events were mapped previously by primer extension (Behler et al. 2018). Transcriptome coverage is given on top for the three analysed strains. Cleavage sites are displayed in the diagrams underneath by black, blue and orange peaks, representing 5'-P (PSS). Since 5'-PPP (TSS) may be converted to 5'-P RNA ends *in vivo* and during RNA-seq library preparation, TSS are partially detected in the PSS signal. Positions which were classified as TSS are labelled with "TSS" next to the respective peaks. For visualization, the average of normalised read counts of the analysed replicates was calculated.

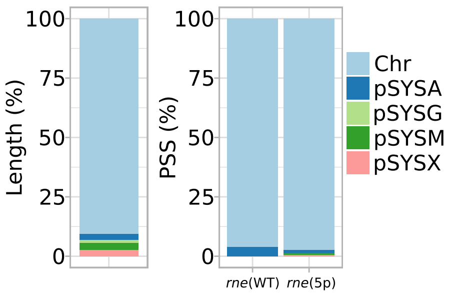


**Supplementary Figure S7.** Relative lengths of different parts of the *Synechocystis* genome (on the left). Percentage of PSS mapped to the five major replicons comparing *rne*(WT) and *rne*(5p) (on the right). Chr: chromosome.

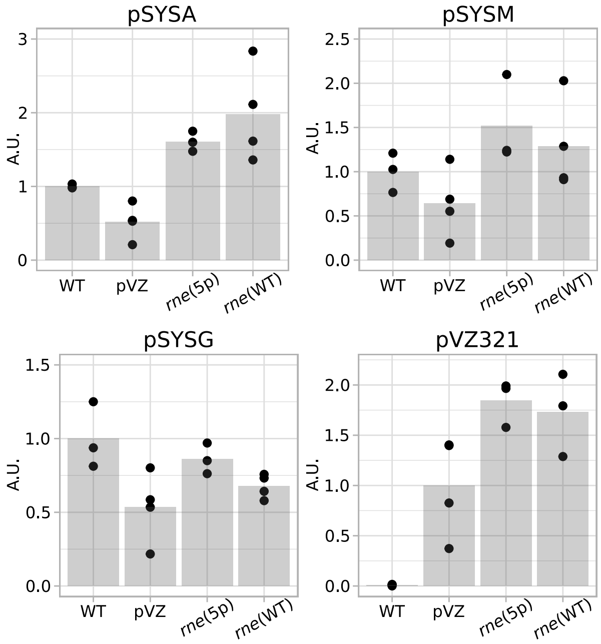


**Supplementary Figure S8.** Quantification of different replicons by qPCR. Data is normalized to signal for a chromosomal locus and to the average detected amount in WT samples except for pVZ321-specific signal for which intensities were normalized to pVZ321 signal and to the average amount detected in the empty-vector control strain pVZ$\Delta$Km$^{\mathrm{R}}$. Black dots represent measurements of biological triplicates, gray bars mean of these. pVZ: pVZ$\Delta$Km$^{\mathrm{R}}$.

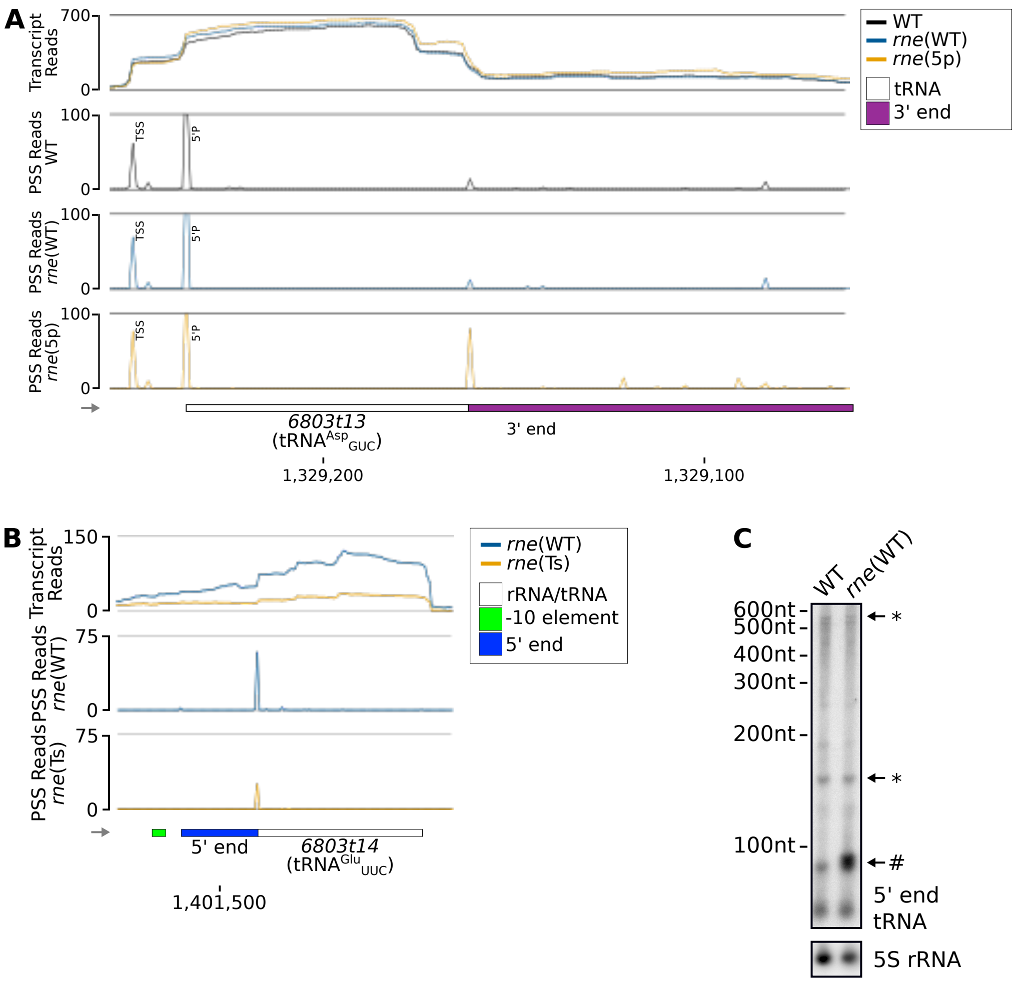


**Supplementary Figure S9.** (**A**) RNA-seq data for tRNA$^{Asp}_{GUC}$ (*6803t13*). Transcriptome coverage is given on top for the three indicated strains. Cleavage sites are displayed in diagrams below by black, blue and orange peaks, representing 5'-P (PSS) detected in the three different strains. Positions which were classified as TSS are indicated by "TSS" next to the respective peak. "5'P" indicates the peak corresponding to the 5' end of the mature tRNA (average read counts WT: 274.84, *rne*(WT): 229.74, *rne*(5p): 147.25). Transcriptome coverage and cleavage sites (PSS) represent the average of normalised read counts of the investigated replicates. (**B**) TIER-seq data comparing *rne*(WT) to a temperature-sensitive strain *rne*(Ts), in which RNase E was inactivated by incubation at 39°C for one hour (Hoffmann et al. 2021). As in (A), transcriptome coverage is given on top for the two indicated strains. Cleavage sites are displayed in diagrams below by blue and orange, representing 5'P (PSS) detected in the two different strains. (**C**) Northern blot of total RNA of wild-type *Synechocystis* (WT) and *rne*(WT) hybridised with a probe targeting the 5' end of *6803t14* (top, 5' end of tRNA) or 5S rRNA (bottom). Asterisks (*) indicate sizes of likely major precursor transcripts, # a processing intermediate accumulating in *rne*(WT) compared to WT. One representative analysis is shown (n=4).

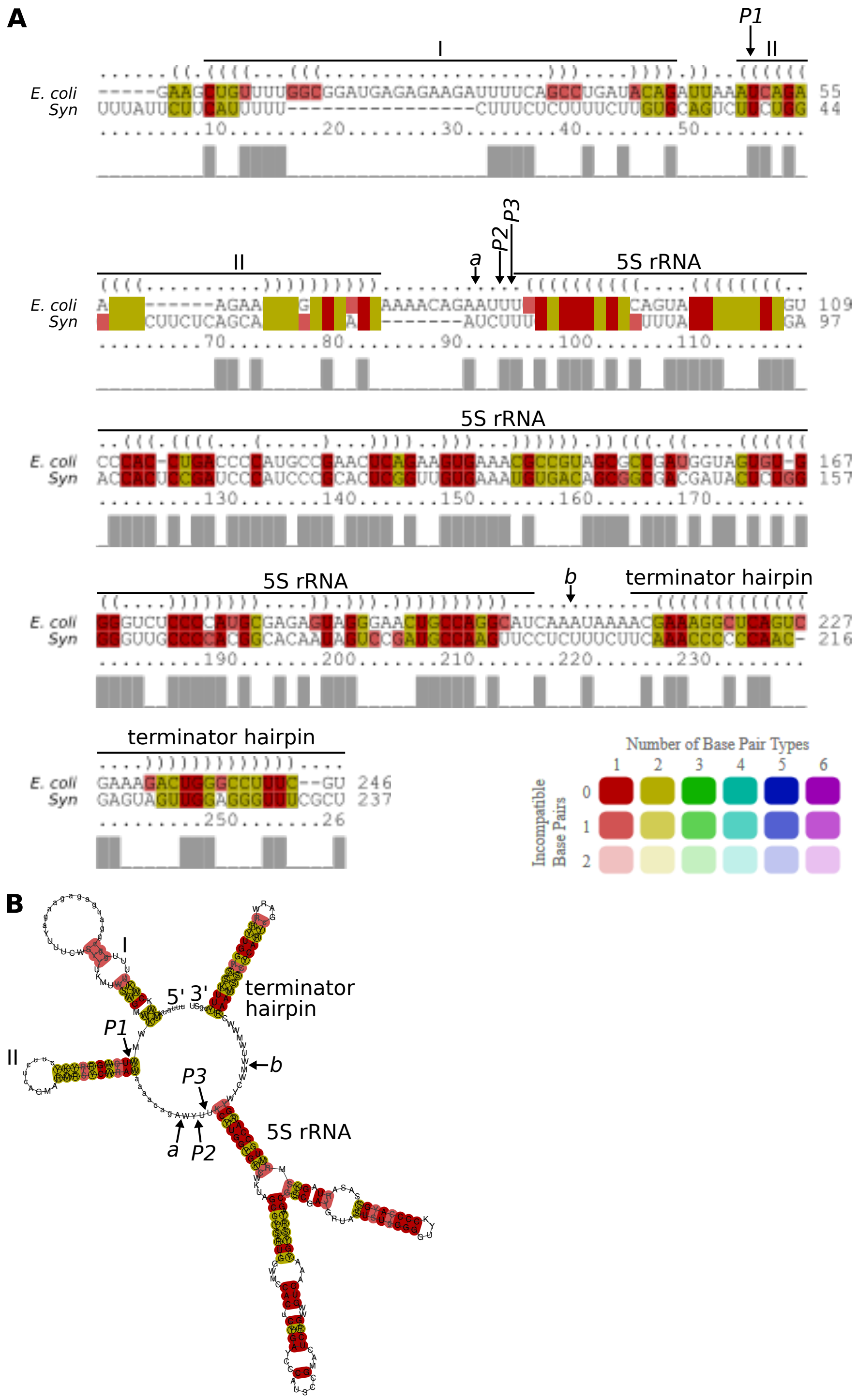


**Supplementary Figure S10.** LocARNA analysis of *E. coli* and *Synechocystis* 9S RNA. (**A**) Sequence alignment of *E. coli* 9S RNA and sequence spanning from 3' end of 23S rRNA to region downstream of 5S rRNA in *Synechocystis* (*Syn*). The end of the aligned sequence of *Synechocystis* RNase E was chosen by investigating read distribution in *rne*(5p) samples. The consensus structure is given above the sequence alignment in dot-bracket notation. (**B**) Consensus structure of both sequences. In panels A and B, colour indicates compatible base pairs. Colour hue indicates the number of compatible base pairing in the corresponding column. Lower colour saturation indicates a higher number of incompatible base pairs. Annotation is based on (Cormack and Mackie 1992). *a* and *b* indicate RNase E cleavage sites mapped in *E. coli* (Basanta, Ghora and Apirion 1978; Cormack and Mackie 1992). *P1*, *P2* and *P3* indicate cleavage positions mapped here.

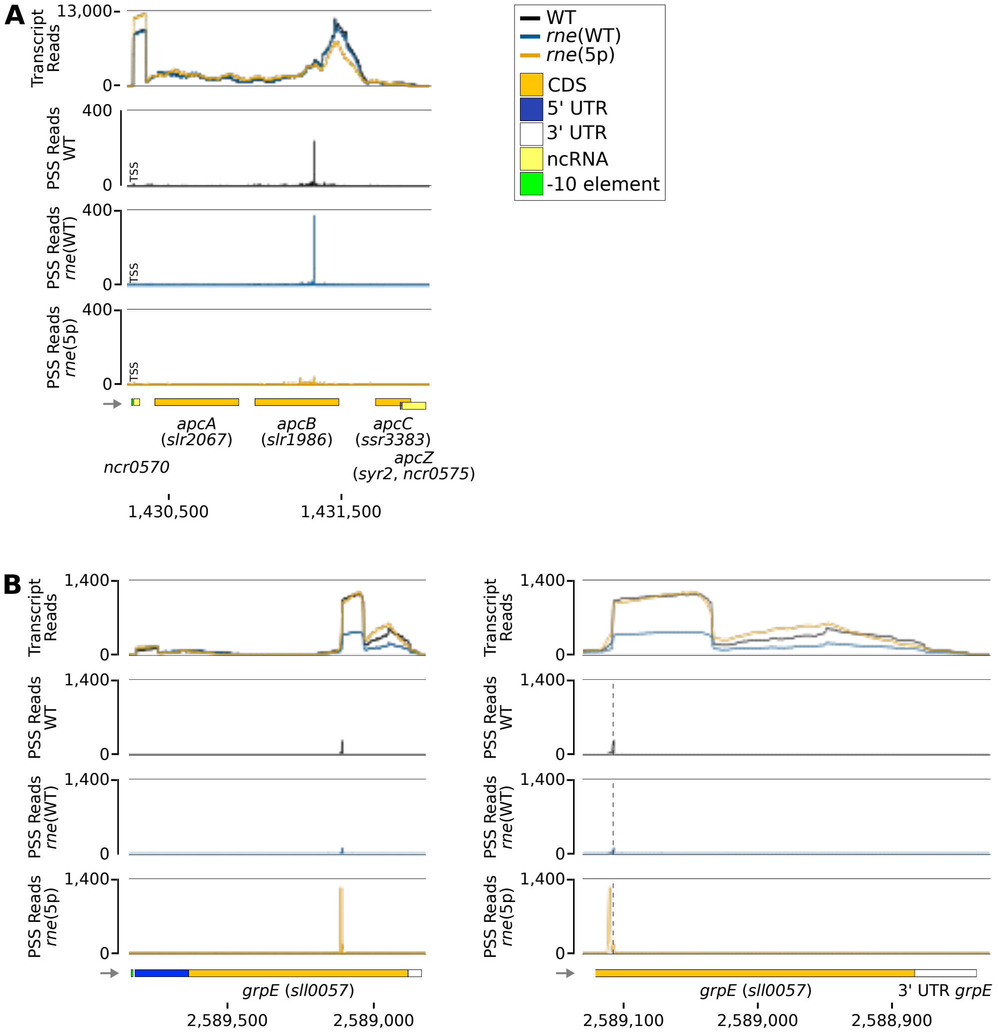


**Supplementary Figure S11.** Examples of 5' sensing dependent cleavage events. (**A**) The *apcABCZ* operon. The gene *apcZ* and the associated -10 element are depicted offset due to spatial constraints. (**B**) Overview of *grpE*. (**C**) Enlarged view of 3' region of CDS of *grpE*. Vertical dashed lines indicate the PSS with most read counts in WT and *rne*(WT). Transcriptome coverage is given on top for the three indicated strains. Cleavage sites are displayed in diagrams below by black, blue and orange peaks, representing 5'-P (PSS) detected in the three different strains. 5'-PPP (TSS) may be converted to 5'-P RNA ends *in vivo* or during RNA-seq library preparation. Thus, TSS are partially detected in the PSS signal. Positions which were classified as TSS are indicated by "TSS" next to the respective peaks. Transcriptome coverage and cleavage sites (PSS) represent the average of normalised read counts of the investigated replicates. CDS: coding sequences, UTR: untranslated region, ncRNA: non-coding RNA.
